# LL-37 modulates IL-17A/F-mediated airway inflammation by selectively suppressing Lipocalin-2

**DOI:** 10.1101/2024.09.03.610924

**Authors:** Anthony Altieri, Dylan Lloyd, Padmanie Ramotar, Anne M van der Does, Mahadevappa Hemshekhar, Neeloffer Mookherjee

## Abstract

**Background:** Levels of the human cationic host defence peptide (CHDP) LL-37 is enhanced in the lungs during neutrophilic airway inflammation. LL-37 drives Th17 differentiation, and Th17 cells produce IL-17A and IL-17F which forms the biologically active heterodimer IL-17A/F. While IL-17 is a critical mediator of neutrophilic airway inflammation, LL-37 exhibits contradictory functions; LL-37 can both promote and mitigate neutrophil recruitment depending on the inflammatory milieu. The impact of LL-37 on IL-17-induced responses in the context of airway inflammation remains largely unknown. Therefore, we examined signaling intermediates and downstream responses mediated by the interplay of IL-17A/F and LL-37, in human bronchial epithelial cells (HBEC). As LL-37 can get citrullinated during airway inflammation, we also examined LL-37-mediated downstream response compared to that with citrullinated LL-37 (citLL- 37), in HBEC.

**Results:** Using an aptamer-based proteomics approach, we identified proteins that are altered in response to IL-17A/F in HBEC. Proteins enhanced in response to IL-17A/F were primarily neutrophil chemoattractants, including chemokines and proteins associated with neutrophil migration such as lipocalin-2 (LCN-2) and Elafin. We showed that selective depletion of LCN-2 mitigated neutrophil migration, thus functionally demonstrating LCN-2 as a critical neutrophil chemoattractant. We further demonstrated that LL-37 and citLL-37 selectively suppresses IL- 17A/F-induced LCN-2 production, in bronchial epithelial cells. Mechanistic studies revealed that LL-37 and citLL-37 suppressed IL-17A/F-mediated C/EBPβ, a transcription factor required for LCN-2 production. In contrast, LL-37 and citLL-37 enhanced the ribonuclease Regnase-1, which is a negative regulator of IL-17 and LCN-2. In an animal model of neutrophilic airway inflammation with elevated IL-17A/F in the lungs, we demonstrated that CRAMP (mouse ortholog of LL-37) negatively correlates with LCN-2.

**Conclusions:** Overall, our findings showed that LL-37 and citLL-37 can selectively suppress the abundance of IL-17A/F-mediated LCN-2, a protein that is critical for neutrophil migration, in bronchial epithelial cells. These results suggest that LL-37, and its modified citrullinated form, has the potential to negatively regulate IL-17-mediated neutrophil migration to control airway inflammation. To our knowledge, this is the first study to report that the immunomodulatory function of LL-37 engages an RNA binding protein, Regnase-1, indicating post-transcriptional regulation of airway inflammation by the peptide.

## Introduction

Cationic Host Defence Peptides (CHDP), also known as antimicrobial peptides, exhibit a wide range of immunity-related functions and selectively regulate inflammatory processes [1–3]. CHDP LL-37 is the only cathelicidin peptide expressed in humans, which influences immune responses in a highly complex manner with at least 16 direct interacting protein partners and altering the expression of over 900 genes with more than a thousand secondary effector proteins [1–3]. LL-37 selectively regulates pathogen- and cytokine-driven inflammation, with mechanisms dependent on the inflammatory environment, and the tissue and cell type [1]. It is known that LL-37 levels are altered in chronic inflammatory diseases of the lung [4, 5]. LL-37 is released from neutrophils by degranulation and NETs (neutrophil extracellular trap) in the lungs during airway inflammation [5]. Previous studies have shown that LL-37 can drive the differentiation of T-helper (Th) cells to Th17 cells in the lungs [6, 7]. Th17 cells produce cytokines IL-17A and IL-17F, and the heterodimer IL-17A/F is a critical pro-inflammatory cytokine that is elevated in neutrophilic airway inflammation during respiratory disease such as severe asthma [8, 9]. Although both LL- 37 and IL-17A/F are elevated in the lungs during airway inflammation, these do not mediate similar effects on neutrophil recruitment and activation. While IL-17 promotes neutrophilic airway inflammation [8–10], the role of LL-37 remains controversial; LL-37 can facilitate recruitment of neutrophils to the lungs to resolve pulmonary infections [11], and in contrast, can limit neutrophil migration and activation in acute lung injury models [12]. The peptide by itself can facilitate neutrophil recruitment and activation [13] but in the presence of an inflammatory mediator it can mitigate neutrophilic inflammation [14]. The impact of LL-37 on IL-17-mediated downstream inflammatory responses and neutrophil recruitment in the lungs remains unresolved.

IL-17A/F targets the IL-17RA/RC receptor complex expressed solely by structural cells such as human bronchial epithelial cells (HBEC), to enhance the recruitment of neutrophils via the production of neutrophil-recruiting chemokines [15–17]. LL-37 can exhibit both pro- and anti-inflammatory functions based on the cellular microenvironment, and the peptide’s interaction with other factors within the inflammatory milieu [18]. A recent study has shown that LL-37 can selectively alter IL-17-mediated pro-inflammatory metabolic and immune responses in synoviocytes [19]. Therefore, in this study we examined the impact of LL-37 on IL-17A/F- mediated responses in HBEC, as both LL-37 and IL-17A/F are enhanced during airway inflammation in the lungs, especially that characterized by neutrophilia..

Recent studies have shown that the arginine residues of LL-37 can get citrullinated by peptidyl arginine deiminase (enzymes enhanced during inflammation), and citrullinated LL-37 (citLL-37) is found in the NETs during airway inflammation [20]. Both LL-37 and citLL-37 are present in the bronchoalveolar lavage fluid of the human lung [20]. Previous studies have shown that citrullination of LL-37 impairs the antimicrobial functions and alters some immunomodulatory functions of the peptide [20–22]. Therefore, in this study we also compared the effects of citLL-37 along with LL-37, in the presence and absence of IL-17A/F in HBEC.

In this study, we used an aptamer-based proteomics array to define proteins altered by IL- 17A/F, in HBEC. We showed that LL-37 and citLL-37 selectively suppress IL-17A/F-induced lipocalin-2 (LCN-2), and that the depletion of LCN-2 mitigates neutrophil migration. We demonstrated that the cathelicidin CRAMP (mouse ortholog of LL-37) negatively correlates with LCN-2 and neutrophil elastase (NE), in an IL-17-driven mouse model of neutrophilic airway inflammation [6]. In mechanistic studies, we showed that LL-37 suppresses IL-17A/F-mediated C/EBPβ, a transcription factor required for LCN-2 production, and in contrast enhances the endoribonuclease Regnase-1 which is an inhibitor of IL-17 and LCN-2 [23]. To our knowledge, this is the first study to demonstrate that LL-37-mediated immunomodulation engages post-transcriptional mechanisms by enhancing the RNA binding protein Reganse-1. Overall, our findings suggest that LL-37 has the potential to selectively limit neutrophilia via the suppression of IL-17A/F-induced LCN-2 production in airway inflammation.

## Materials and Methods

### Reagents

Peptides LL-37 and sLL-37 were manufactured by CPC Scientific (Sunnyvale, CA, USA), and citLL-37 was obtained from Innovagen AB (Lund, Sweden). Table 1 shows the sequences of the peptides used. Peptides reconstituted in endotoxin-free E-Toxate^TM^ water were aliquoted and stored in glass vials at -20°C and used within 3 months of reconstitution. Peptides were thawed at room temperature (RT), sonicated for 30 seconds, and vortexed for 15 seconds before use. Recombinant human cytokines IL-17A/F (carrier free) was obtained from R&D Systems (Oakville, ON, CA).

**Table 1.**
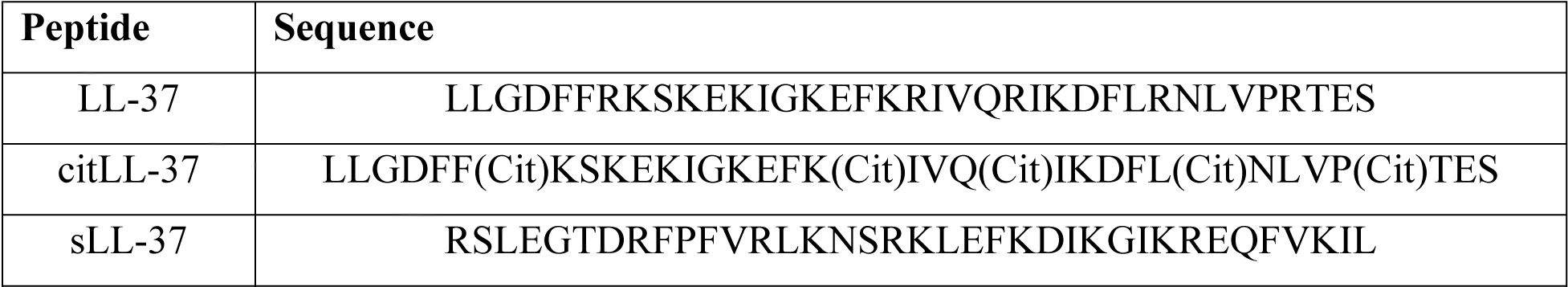
Peptide Sequences.

### Cell culture

HBEC-3KT cells (ATCC® CRL-4051™) were cultured using airway epithelial cell basal medium (ATCC® PCS-300-030™) supplemented with bronchial epithelial cell growth kit (ATCC® PCS-300-040™), as previously described [24]. Cell culture medium was changed to airway epithelial cells basal medium containing 6 mM L-glutamine without growth factors, 24 h prior to stimulation with various cytokines as indicated. Human primary bronchial epithelial cells (PBEC) were expanded for biobanking as previously described [25–27]. PBECs were isolated from resected tumor-free lung tissues obtained from four anonymized donors. Cells were isolated from macroscopically normal lung tissue obtained from patients undergoing resection surgery for lung cancer at the Leiden University Medical Center (LUMC), the Netherlands. These patients were enrolled via a no-objection system for coded anonymous further use of such tissue (www.coreon.org). However, since September 2022, patients were enrolled using active informed consent in accordance with local regulations from the LUMC biobank with approval by the institutional medical ethical committee (B20.042/Ab/ab and B20.042/Kb/kb). After thawing, PBECs were expanded in T75 flasks pre-coated with coating media (containing 30 µg/mL PureCol (Advanced Biomatrix, California, USA), 10 µg/mL fibronectin (Sigma), 10 µg/mL BSA (Sigma) in PBS (Gibco)), in supplemented keratinocyte serum-free medium (Gibco) containing 0.2 ng/mL epidermal growth factor (Life Technologies), 25 µg/mL bovine pituitary extract (Gibco), 1 µM isoproterenol (Sigma) and 1:100 dilution of antibiotics Penicillin and Streptomycin (Lonza), and maintained until ∼ 80% confluent. PBECs were seeded (5000/cm^2^) tissue culture (TC) plates pre-coated with coating media (detailed above), and cultured with a 1:1 mixture of supplemented Dulbecco’s modified Eagle’s medium (Gibco) with a 1:40 dilution of HEPES (Invitrogen), and basal bronchial epithelial cell medium (ScienCell) containing bronchial epithelial cell growth supplement (ScienCell), 1:100 dilution of Penicillin/Streptomycin and 50 nM of a light stable analog of retinoic acid, EC-23 (Tocris, UK). The culture medium was replaced every 48 h, and the medium was replaced with medium without EGF, BPE, BSA and hydrocortisone (starvation media) 24 h prior to stimulation with cytokine and/or peptides.

### ELISA

Tissue culture (TC) supernatants were centrifuged (250xg at RT for 5 minutes) and the cell-free TC supernatants were aliquoted, stored at -20°C until use. The abundance of LCN2, Elafin, GROα and CCL20 were measured in the TC supernatants using related ELISA kits obtained from R&D Systems, as per the manufacturer’s instructions.

### Slow Off-rate Modified Aptamer (SOMAmer)-based proteomic array

HBEC-3KT cells were stimulated with IL-17A/F (50 ng/mL) for 24 h. Total cell lysates were obtained with lysis buffer (M-PER™ and HALT protease and phosphatase inhibitor cocktail obtained from ThermoFisher Scientific, Burlington, ON, Canada) and protein concentration was determined in each sample by microBCA protein assay (Thermo Fisher Scientific). Cell lysates (14 µg of total protein each) obtained from from five independent experiments were probed independently using the Slow off-rate Modified Aptamer (SOMAmer^®^) V.2 proteomic array, as previously described by us [17, 24]. The SOMAmer® V.2 protein arrays profiled the abundance of 1322 protein targets in each sample as detailed in previous studies [17, 28–31]. Protein abundance was quantified using the Agilent hybridization array scanner in relative fluorescence unit (RFU). The RFU readout values were log2-transformed and used for pairwise differential analysis as indicated in individual figure legends. Heatmap with hierarchical clustering was generated using the Multi-Experiment Viewer Version 10.2 and GraphPad PRISM 9 was used for visual representation of changes in protein expression profile.

### Quantitative Real-Time PCR (qRT-PCR)

Total RNA was isolated from cells using the MagMax^Tm^-96 Total RNA isolation kit according to the manufacturer’s instructions. mRNA abundance was assessed by the SuperScript III Platinum Two-Step qRT-PCR Kit with SYBR Green (Invitrogen) according to the manufacturer’s instructions, using the QuantStudio 3 (Applied Biosystems, CA, USA) machine, as previously described by us [32]. Quantitect Primer Assays (Qiagen) were used for detection of *ARID5A* (#QT00049672), *ZCH312A* (#QT00229838), *NFKBIZ* (#QT00049672), *CEBPB* (#QT00237580) and *18S RNA* (#QT00199367). Relative fold changes were calculated using the comparative ΔΔCt method [33] after normalization with 18S RNA as the reference gene, as previously described by us [24].

### Western blots

HBEC-3KT cells were washed with cold PBS, scraped using a 25 cm cell scrapper (VWR) and collected in PBS containing 1X protease inhibitor cocktail (Cell Signaling Technology, Massachusetts, USA). Cells were lysed in cold PBS containing 1X Cell Lysis Buffer (Cell Signaling Technology) containing protein inhibitor cocktail (New England Biolabs). Cell lysates were incubated on ice for 5 minutes, sonicated in a water bath sonicator for 15 seconds, and centrifuged at 14,000xg at 4°C for 10 minutes to obtain cell-free lysates. Total protein concentration was determined in each lysate using microBCA protein assay (Thermo Fisher Scientific). Cell lysates (25 μg total protein) were resolved on 4-12% NuPage^TM^ 10% Bis-Tris Gels (Invitrogen) followed by transfer to nitrocellulose membranes (Millipore, Massachusetts, USA). Membranes were blocked with Tris-buffered saline (TBST) (20 mM Tris–HCl, pH 7.5, 150 mM NaCl, 0.1% Tween-20) containing 5% Bovine Serum Albumin (BSA) and probed with different primary antibodies, and secondary antibodies for detection, in TBST containing 2.5% BSA. Primary antibodies specific to human MCPIP1/Regnase-1 was obtained from Abcam (Toronto, ON, Canada). Antibodies specific to human NF-κB p65, phospho-IKKα/β, IκB-ζ, and C/EBPβ were obtained from Cell Signaling Technology (New England Biolabs, ON, Canada). The antibody specific to human Arid5a was obtained from Sigma (Toronto, ON, Canada). Anti-human β-actin antibody was obtained from Millipore (Burlington, MA, USA). HRP-linked purified anti-rabbit IgG- and anti-mouse IgG- secondary antibodies were obtained from Cell Signaling Technology. HRP-linked purified anti-goat IgG secondary antibody was obtained from Abcam (Toronto, ON, Canada). The blots were developed using ECL Prime detection system (Thermo Fisher Scientific) according to the manufacturer’s instructions.

### Neutrophil isolation and migration

Neutrophil isolation was performed as previously described by us [24]. Briefly, venous blood was collected in EDTA vacutainer tubes from healthy volunteers with written informed consent according to a protocol approved by the University of Manitoba Research Ethics Board. Human neutrophils were isolated using the EasySep^TM^ Direct Human Neutrophil Isolation Kit (STEMCELL technologies Canada Inc., Vancouver, BC, Canada), according to the manufacturer’s protocol, using ∼25 mL of blood with the isolation cocktail containing 50 µL RapidSpheres^TM^ provided in the kit, to obtain enriched human neutrophils by negative selection. Neutrophils isolated from human blood (6 x 10^5^ cells/well, 200 µL total volume) were added to the upper chamber of the inserts on 5 µM permeable polycarbonate membrane Transwell supports (Costar, Corning, NY, USA) for neutrophil migration assay as follows.

TC supernatants (600 µL) were collected from HBEC-3KT cells stimulated with IL-17A/F (50 ng/mL) and TNFα (20 ng/mL) after 24 h, added to the bottom chamber of the Transwell plates. Plates were incubated at 37°C in a humidified chamber with 5% CO2 for 30 min, prior to addition of the human neutrophils to the upper chamber. Human recombinant neutrophil cytokine IL-8 (30 ng/mL) in the airway epithelial cells basal medium (containing 6 mM L-glutamine) was used in the bottom chamber as a positive control [24]. Subsequently the Transwell plates were incubated for 2 h, and the number of neutrophils that migrated to the bottom chamber was counted using a Scepter^TM^ 2.0 Handheld Automated Cell Counter (Millipore Ltd, ON, Canada).

### Immunodepletion of LCN-2

LCN-2 was depleted from TC supernatants as indicated, using an immunodepletion approach with a Dynabeads^TM^ Protein G Immunoprecipitation Kit (Invitrogen), according to the manufacturer’s instructions. Briefly, the magnetic beads were incubated with an anti-LCN2 (Abcam) or IgG isotype control (Abcam) antibody, for 2 h at room temperature (RT). The TC supernatants were incubated with either anti-LCN2 or IgG bound magnetic beads as indicated, for 2 h at RT. The resulting TC supernatants (600 uL) were added to the bottom chamber of a Transwell TC plate for assessing neutrophil migration as detailed above.

### Mouse model of neutrophilic airway inflammation

The mouse model used in this study was previously demonstrated to induce a IL-17-dependent neutrophilic airway inflammation by co-sensitization with allergen house dust mite (HDM) and low dose endotoxin, followed by allergen (HDM) only recall challenge, using only male mice [6]. Therefore, in this study we have optimized this model for both male and female mice, and we provide data using sex-disaggregated data analysis to demonstrate sex-related differences in outcomes assessed. Briefly, male and female BALB/c mice (6 to 7 weeks) were obtained from Charles River Laboratories, randomly sorted within sexes, and housed maximum 5 mice per cage. The mice were acclimatized for one week and subsequently sensitized with intranasal (i.n) administration of HDM extract (∼25 µg per mouse) with low dose lipopolysaccharide (1 µg per mouse), once daily for three days. Mice were further rested for 4 days and subsequently challenged with HDM alone (allergen-recall phase), once daily for 8 days. The HDM protein extract used was of low endotoxin content i.e. <300 EU/mg protein weight [34] (Greer Laboratories, Lenoir, NC, USA). Control groups with either saline alone or LPS alone during the sensitization phase were subsequently challenged with saline in the allergen recall phase. Mice were anesthetized using sodium pentobarbital (90 mg/kg) 24 h after the last HDM challenge, followed by tracheostomy, the lung was washed twice with 1 mL of cold saline to obtain bronchoalveolar lavage (BAL) samples. Lung tissues were obtained from the right lung middle lobe and collected in Tissue Protein Extraction Reagent (Pierce; ThermoFisher Scientific, Rockford, IL, USA) containing 1X Protease Inhibitor Cocktail (Sigma Aldrich, Oakville, ON, Canada). The animal model protocol used in this study was approved by the University of Manitoba Animal Research Ethics Board (protocol #18-038), and is compliant with ARRIVE guidelines for *in vivo* animal research [35].

### Cell differential assessment

BALF obtained was centrifuged (150xg at RT for 10 minutes) and the cell pellet was resuspended in 1 mL sterile saline. Cell suspension (100 µL) in a Cytospin slide stained with a modified Wright–Giemsa stain (Hema 3® Stat Pack, Fisher Scientific, Hampton, NH, USA) was used for cell differential counts using a Carl Zeiss Axio Lab A1 (Carl Zeiss, Oberkochen, Germany) microscope for imaging. Cells were counted blinded by two different individuals in 8-10 image frames at 20X magnification per slide.

### Cytokine detection in bronchoalveolar lavage fluid (BALF) and lung tissues

BALF samples were centrifuged (150xg for 10 minutes at 4°C) to obtain cell-free supernatants. Lung tissues were homogenized on ice using the Cole-Parmer LabGEN 125 Homogenizer (Canada Inc, Montreal, QC, Canada). Lung tissue homogenates were centrifuged (10,000xg at 4°C) to obtain lung tissue lysates, and total protein abundance in the lysates was quantified using a Bicinchoninic acid (BCA) Protein Assay (Pierce). BALF and lung tissue lysates were aliquoted and stored at -20°C and - 80°C respectively until use. Abundance of the mouse cathelicidin peptide CRAMP was measured by ELISA (Biomatik, Kitchener, Ontario, Canada). Abundance of LCN-2 and IL-17A/F were measured by Quantikine ELISA assays from R&D Systems (Minneapolis, MN, USA). BALF (50 μL) was used to assess the abundance of LCN-2 and IL-17A/F, and 100 µL of BALF was used to determine the abundance of CRAMP. Lung tissue lysates with 50 µg of total protein was used for cytokine evaluation.

### Statistical analyses

Specific statistical analyses used are detailed in each figure legend. Pairwise differential analysis was conducted on normalized log2 protein expression values, and Welch’s t-test with a cutoff of *p<0.05* was used to select protein abundance changes that were significantly altered in response to IL-17A/F in the high-content aptamer-based proteomic array. Repeated measures one-way ANOVA with Fisher’s least significant difference test was used for statistical analysis was used to determine proteins that were enhanced in TC supernatant or cell lysate in response to IL-17A/F, and altered in response to LL-37, citLL-37, and sLL-37 in the presence/absence of IL-17A/F in HBEC-3KT and PBEC. In addition, repeated measures one-way ANOVA with Fisher’s least significant difference test was used for statistical analysis was used to determine changes to mRNA abundance in response to LL-37, citLL-37, and sLL-37 in the presence/absence of IL-17A/F in HBEC-3KT. One-way ANOVA with Fisher’s LSD test was used to determine changes in cell accumulation as well as protein targets in the lungs in the mouse model of neutrophilic airway inflammation. Pearson’s correlation analysis was performed to determine the correlations between select protein targets in the lungs of the mouse model of neutrophilic airway inflammation.

## Results

### IL-17A/F alters the bronchial epithelial proteome and significantly increases the abundance of neutrophil chemotactic proteins

HBEC-3KT cells were stimulated with IL-17A/F (50 ng/mL) for 24 h, and total cell lysates (14 µg of total protein each) obtained from five independent experiments were probed independently using the Slow off-rate Modified Aptamer (SOMAmer^®^) V.2 proteomic array. To determine overall changes in the bronchial epithelial proteome in response to IL-17A/F, a pairwise differential analysis was performed on normalized log2 protein abundance values of HBEC-3KT stimulated IL-17A/F, compared to unstimulated cells. IL-17A/F significantly (p<0.05) altered the abundance of 25 proteins (Supplementary Table I), increasing the abundance of 20 and suppressing the abundance of 5 proteins, compared to unstimulated cells (Figure 1A). Proteins that were enhanced by >2-fold (p<0.05) in response to IL-17A/F, were predominantly neutrophil chemotactic factors. The two most enhanced proteins following stimulation with IL-17A/F were CHDPs LCN-2 and Elafin. LCN2 is also shown to promote neutrophil recruitment [36, 37]. Elafin is known to prevent damage to the airway epithelium during lung inflammation by degrading serine proteases such as NE [38, 39]. In addition, neutrophil-recruiting chemokine GROα was also significantly enhanced by IL-17A/F [40].

**Figure 1:**
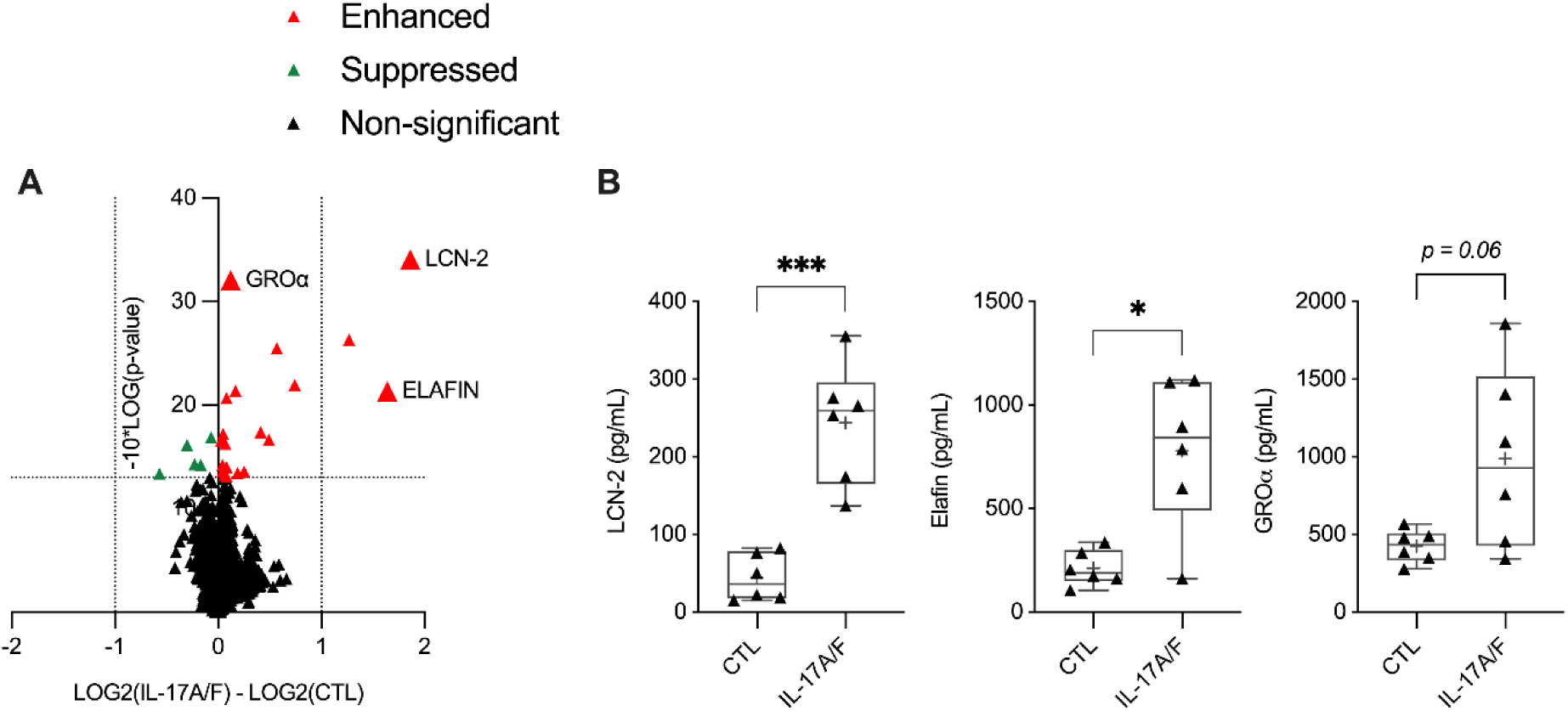
IL-17A/F alters the bronchial epithelial cell proteome. HBEC-3KT cells were stimulated with IL-17A/F (50 ng/mL) for 24 h. **(A)** Cell lysates from cells stimulated with IL- 17A/F and unstimulated cells (14 µg total protein per sample), obtained from five independent experiments (n=5), were independently probed using the high-content aptamer-based proteomic array. Pairwise differential analysis was conducted on normalized log2 protein expression values, and Welch’s t-test with a cutoff of *p<0.05* was used to select protein abundance changes that were significantly altered in response to IL-17A/F. **(B)** TC supernatant collected from cells 24 h post- stimulation was examined for the abundance of LCN-2, Elafin, and GROα, by ELISA. Each dot represents an independent experiment (n=5), and bars show the median and min-max range. Repeated measures one-way ANOVA with Fisher’s least significant difference test was used for statistical analysis (*\*p≤0.05, ***p≤0.005*).

As secreted proteins primarily mediate cellular communication, we further examined the abundance of these three selected proteins i.e. LCN-2, Elafin and GROα in TC supernatants by ELISA, in independent experiments. Consistent with the results obtained from the proteomic profiling of cell lysates (Figure 1A), the abundance of LCN-2, Elafin and GROα were enhanced in the TC supernatants of HBEC-3KT cells stimulated with IL-17A/F, after 24 h (Figure 1B). Therefore, these protein candidates (LCN-2, Elafin and GROα) were selected to examine the effect of CHDP LL-37 and its citrullinated form citLL-37, on IL-17A/F-mediated responses in bronchial epithelial cells in further studies.

### LL-37, and citrullinated LL-37, suppresses IL-17A/F-mediated LCN-2 production in bronchial epithelial cells

As discussed above, although both LL-37 and IL-17A/F are enhanced in the lungs during neutrophilic airway inflammation, the impact of the interplay of these molecules on neutrophil recruitment remains unresolved. Previous studies have demonstrated that LL-37 indirectly promotes neutrophil recruitment by inducing chemokine secretion [32]. However, it has also been shown that in presence of an inflammatory mediator LL-37 can selectively mitigate neutrophilic inflammation [14]. Therefore, we examined the impact of LL-37 on IL-17A/F- mediated production of neutrophil-recruiting proteins selected from the aptamer-based proteomics array data (Figure 1), in bronchial epithelial cells. As the concentration of LL-37 in individuals with neutrophilic airway inflammation ranges from 0.25 μM to 1 μM [5], we first performed dose titrations of LL-37 to select a concentration within this physiological range that results in enhancement of chemokines by the peptide alone, without eliciting cytotoxicity as demonstrated in our previous study [41]. HBEC-3KT cells were stimulated with LL-37 and scrambled LL-37 (sLL-37) as a negative control, for 24 h. The abundance of chemokines GROα and IL-8 was measured in TC supernatants by ELISA. LL-37 at both 0.25 μM and 0.5 μM concentrations increased the abundance of GROα and IL-8 (Supplemental Figure 1). Based on these results, we selected 0.25 μM for further studies.

It is known that LL-37 is citrullinated, which is a post-translational modification wherein arginine residues are converted to citrulline, in the lungs during neutrophilic airway inflammation [20–22, 42, 43]. Therefore, we performed a comparative analysis of the effects of LL-37 and citrullinated LL-37 (citLL-37) in this study. To determine the effects of the peptides by itself, HBEC-3KT cells were stimulated with 0.25 μM of peptides LL-37, citLL-37, or sLL-37, and the abundance of chemokines GROα and IL-8 was measured in TC supernatants by ELISA after 24 h. LL-37 significantly increased the abundance of GROα (by ∼26%) compared to unstimulated cells, however citLL-37 did not (Supplemental Figure 2). Although both LL-37 and citLL-37 significantly increased the abundance of IL-8 (Supplemental Figure 2), increase in IL-8 in response to citLL-37 was ∼33% less (*p=0.07*) compared to that enhanced by LL-37.

To examine the effect of LL-37 and citLL-37 on IL-17A/F-mediated response, HBEC-3KT were stimulated with 0.25 μM of LL-37, citLL-37, or sLL-37, in the presence and absence of IL- 17A/F (50 ng/mL) for 24 h. TC supernatants were used to examine the abundance of proteins selected from the proteomic array (Figure 1), namely LCN-2, Elafin and GROα, by ELISA. Both LL-37 and citLL-37 significantly enhanced GROα production in the presence of IL-17A/F by ∼300% and ∼210% respectively, compared to control (Figure 2). Thus, the increase in GROα was less in the presence of citLL-37 compared to that mediated by LL-37 by ∼100%. The peptides did not alter IL-17A/F-mediated Elafin production (Figure 2). In contrast, both LL-37 and citLL-37 suppressed IL-17A/F-mediated LCN-2 production by ∼50% in HBEC-3KT (Figure 2). These results indicated that LL-37 and citLL-37 enhance IL-17A/F-mediated chemokine GROα, and in contrast suppress LCN-2.

**Figure 2:**
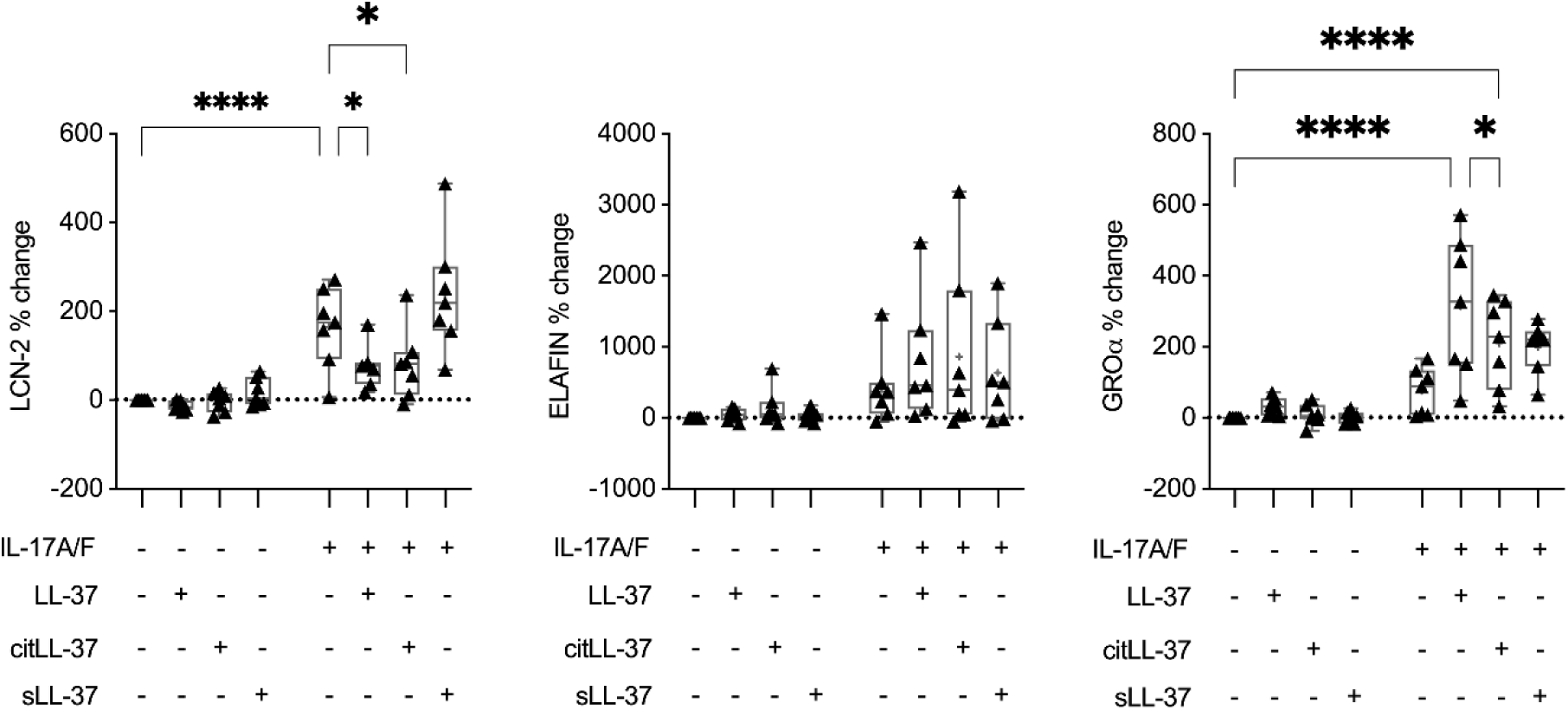
LL-37 and citLL-37 selectively alter IL-17A/F-mediated protein production in bronchial epithelial cells. HBEC-3KT cells were stimulated with either LL-37, citLL-37 or sLL- 37 (0.25 μM), in the presence and absence of IL-17A/F (50 ng/mL), for 24 h. TC supernatants collected from seven independent experiments (n=7) were examined by ELISA for the abundance LCN-2, Elafin and GROα. Y-axis represents % change compared to paired unstimulated controls. Each dot represents an independent experiment, and bars show the median and min-max range. Each dot is reported as percent (%) change compared to unstimulated cells where % change = ((treatment – control) / control) x 100%. Repeated measures one-way ANOVA with Fisher’s least significant difference test was used for statistical analysis (*\*p≤0.05, ****p≤0.0001*).

### Depletion of LCN-2 mitigates neutrophil migration

As LL-37 and citLL-37 suppresses IL- 17A/F-induced LCN-2 in HBEC (Figure 2), it is likely that these peptides can intervene in neutrophil migration to the lungs. Although previous studies suggest a neutrophil chemoattractant function for LCN-2 [24], there is no direct evidence demonstrating LCN-2’s ability to promote neutrophil migration in the context of airway inflammation. Therefore, we performed a functional neutrophil migration assay to determine the direct effect of LCN-2 on neutrophil migration, in the presence and absence of an anti-LCN-2 antibody. We have previously shown that the combination IL-17A/F and TNFα is required for neutrophil migration, and that this cytomix (IL-17A/F+ TNFα) robustly enhances both LCN-2 and GROα production in bronchial epithelial cells, compared to either cytokine alone [24]. Based on our previous study [24], HBEC-3KT cells were stimulated with the combination of IL-17A/F (50 ng/mL) and TNFα (for 20 ng/mL) for 24 h. TC supernatants were incubated with an anti-LCN-2 antibody to deplete LCN-2, and the abundance of LCN-2 was examined by ELISA to assess the efficiency of LCN-2 immunodepletion. Aligned with our previous results [24], the cytomix IL-17A/F+TNFα robustly enhanced LCN-2 abundance in TC supernatant, and LCN-2 was significantly reduced in the TC supernatants treated with the anti-LCN-2 antibody (Supplemental Figure 3). These TC supernatants, with and without LCN-2 immunodepletion, were used in the bottom of a Transwell plates for a neutrophil migration assay, as previously described by us [24]. Recombinant chemokine IL-8 (30 ng/mL) was used as a positive control.

TC supernatants collected from cells treated with the cytomix (IL-17A/F+TNFα) significantly enhanced neutrophil migration (Figure 3), corroborating our previous study [24]. There was no difference in neutrophil migration mediated by TC supernatants with and without the IgG isotype control (Figure 3). Whereas, TC supernatants after immunodepletion of LCN-2 resulted in a significant reduction in IL-17A/F+TNFα-mediated neutrophil migration, compared to that mediated by TC supernatants with and without IgG isotype control (Figure 3). These results demonstrated that neutrophil migration mediated by factors secreted from bronchial epithelial cells is dependent on LCN-2. As LL-37 and citLL-37 suppresses LCN-2 in bronchial epithelial cells (Figure 2), and LCN-2 is critical for neutrophil migration (Figure 3), our results taken together suggest that the peptides have the potential to limit neutrophil infiltration in the lungs.

**Figure 3:**
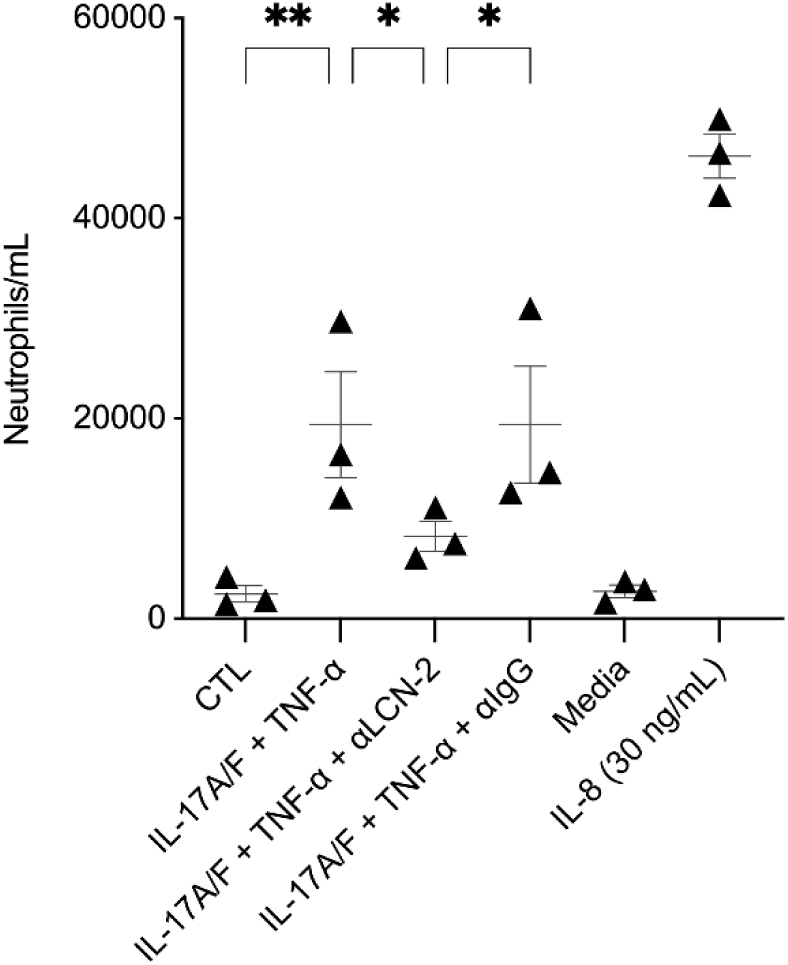
Antibody-mediated neutralization of LCN-2 suppresses neutrophil migration. HBEC- 3KT cells were stimulated with IL-17A/F (50 ng/mL) + TNFα (20 ng/mL) for 24 h. TC supernatants were collected and used in the bottom chamber of transwell plates in a neutrophil migration assays, wherein neutrophils isolated from human blood were used in the upper chamber of the transwell plates. Cell culture medium spiked with human recombinant IL-8 (30 ng/mL) was used as a positive control for neutrophil migration. Results are shown as boxplots with the median line and IQR, and whiskers show minimum and maximum values. Each data point represents an independent experimental replicate with TC supernatant (n=3), using neutrophil isolated from one donor. Each dot represents the average number of neutrophils that traversed the membrane within two hours in each experiment, and bars show the mean and SEM. Repeated measures one-way ANOVA with Fisher’s least significant difference test was used for statistical analysis (*\*p≤0.05, **p≤0.01*).

### LL-37 and citLL-37 disparately alters the mRNA abundance of transcription factors NFKBIZ and CEBPB

Previous studies have demonstrated that IL-17A engages disparate transcription factors (TF) to regulate the production LCN-2 and GROα in epithelial cells [44, 45]. IL-17A- mediated induction of TF IκBζ and C/EBPβ are both required for the transcription of LCN-2, whereas the transcription of GROα is solely dependent on IκBζ [44, 45]. As we have shown that LL-37 and citLL-37 disparately changes IL-17A/F-mediated production of LCN-2 and GROα (Figure 2), we further examined the transcriptional response of *NFKBIZ* (which encodes IκB-ζ) and *CEBPB* (which encodes C/EBPβ), in response to the peptides in bronchial epithelial cells.

HBEC-3KT were stimulated with LL-37, citLL-37, or sLL-37 (0.25 μM), in the presence and absence of IL-17A/F (50 ng/mL), and the mRNA abundance of *NFKBIZ* and *CEBPB* were examined by qRT-PCR, 3 and 6 h post-stimulation. IL-17A/F stimulation significantly enhanced the mRNA abundance of *NFKBIZ*, >2-fold after 3 h and ∼5-fold after 6 h, compared to unstimulated cells (Figure 4). IL-17A/F-mediated increase in *NFKBIZ* mRNA was further enhanced by ∼3-fold by LL-37, but not significantly by citLL-37, after 3 h (Figure 4A). However, at the later time point of 6 h post-stimulation, IL-17A/F-mediated increase in *NFKBIZ* mRNA was not altered by the peptides (Figure 4B). In response to IL-17A/F, mRNA abundance of *CEBPB* was significantly enhanced ∼1.5-fold compared to unstimulated cells (Figure 4B), and this was significantly suppressed in the presence of both LL-37 and citLL-37 to baseline, 6 h post-stimulation (Figure 4B). Taken together, these results indicate that the IL-17A/F-mediated increase in IκB-ζ, which is required for the transcription of GROα, is significantly enhanced by LL-37 but not as robustly by citLL37. In contrast, both LL-37 and citLL-37 suppresses IL-17A/F-mediated enhancement of C/EBPβ, a TF required for LCN-2 production. These results demonstrate a differential regulation of TF by LL37 and citLL-37 to modulate IL-17A/F response in HBEC.

**Figure 4:**
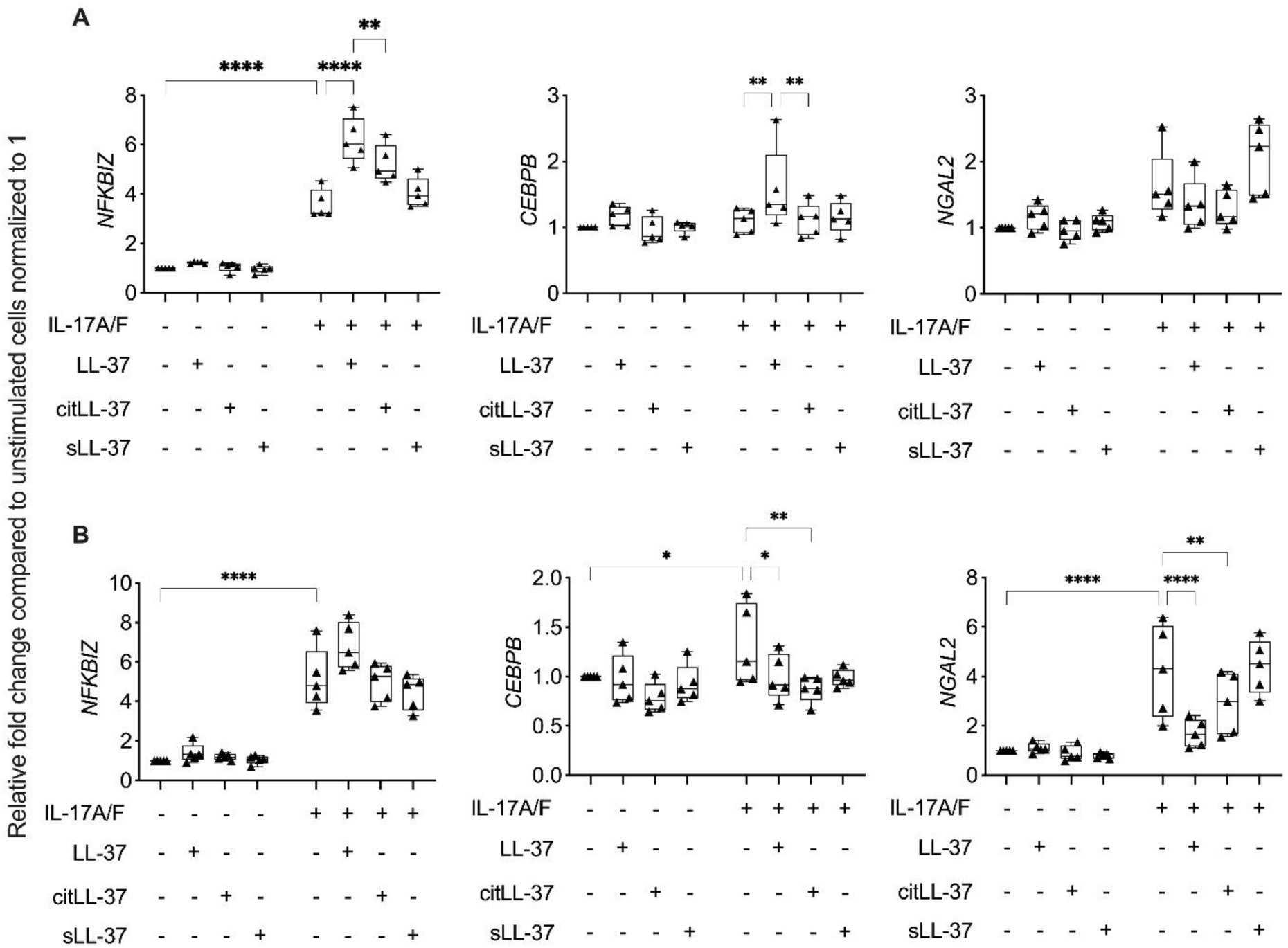
Differential effect of LL-37 and citLL-37 on IL-17A/F-mediated NFKBIZ and CEBPB mRNA abundance. HBEC-3KT cells (n=5) were stimulated with either LL-37, citLL-37 or sLL-37 (0.25 μM), in the presence and absence of IL-17A/F (50 ng/mL). mRNA was isolated after **(A)** 3 h and **(B)** 6 h, and the mRNA abundance of *NGAL2*, *NFKBIZ* and *CEBPB* was examined by qRT-PCR. Fold changes (Y-axis) for each gene candidate was normalized to 18S RNA, and compared to unstimulated cells normalized to 1, using the comparative ΔΔCt method. Each data point represents an independent experimental replicate and bars show the mean and SEM. Fisher’s LSD test for one-way ANOVA was used to determine statistical significance (*\*p<0.05, **p<0.01, ***p<0.001, ****p<0.0001*).

As the mRNA abundance of C/EBPβ, one of the TFs required for the transcription of LCN-2, is suppressed by both LL-37 and citLL-37 (Figure 4B), we also examined the mRNA abundance of *NGAL2* (gene for LCN-2). IL-17A/F-mediated increase in *NGAL2* mRNA abundance was significantly suppressed by both LL-37 and citLL-37 (to baseline levels), after 6 h (Figure 4B). This is consistent with the protein data, wherein IL-17A/F-mediated LCN-2 production is significantly suppressed by both the peptides (Figure 2). As both LL-37 and citLL-37 significantly suppressed IL-17A/F-mediated enhancement of the TF C/EBPβ and the mRNA abundance of *NGAL2* (Figure 4B), these results suggest that a mechanism related to the suppression of IL-17A/F- mediated LCN-2 production by these peptides is by intervening in C/EBPβ-induced signaling cascade.

### LL-37 and citLL-37 enhances the RNA binding protein Regnase-1

Previous studies have shown that IL-17 signaling cascade activates competing RNA binding proteins (RBP), Arid5a and endoribonuclease Regnase-1 (also known as MCPIP1) to regulate its downstream endpoints including LCN-2. Arid5a promotes IL-17-mediated downstream responses, and in contrast Regnase-1 is a negative regulator inhibiting IL-17-mediated responses [23, 44]. Therefore, we examined the effect of LL-37 and citLL-37 on these RBP. HBEC-3KT cells were stimulated with IL-17A/F (50 ng/mL) in the presence and absence of the peptides LL-37, citLL-37 or sLL-37 (0.25 μM each), and the abundance of Arid5a and Regnase-1 was examined by Western blots, after 30 minutes and 24 h. LL-37 and citLL-37 enhanced Arid5a protein abundance, and IL-17A/F- mediated increase in Arid5a was further enhanced by both the peptides, after 30 min (Supplemental Figure 4A). However, at the later time point of 24 h post-stimulation, only LL-37 maintained enhancement of Arid5a abundance (*p*<0.06), and IL-17A/F-mediated changes in Arid5a were not altered by the peptides (Supplemental Figure 4B). In contrast, the negative regulator Regase-1 was significantly enhanced by both LL-37 and citLL-37 compared to unstimulated cells (Supplemental Figure 4A). Representative Western blot images are provided in Supplemental Figure 4C. To our knowledge this is the first study to report LL-37-mediated increase of a RBP in the context of a CHDP-mediated immunomodulatory functions. As this is a novel finding in LL-37 immunobiology, we further examined the abundance of Regnase-1 and LCN-2, and associated signaling intermediates, in human primary bronchial epithelial cells (PBEC).

Human PBEC were stimulated with LL-37, citLL-37, or sLL-37 (0.25 μM), in the presence and absence of IL-17A/F (50 ng/mL). The abundance of Regnase-1 was examined by Western blots after 30 min and 24 h (representative Western blots are shown in Supplemental Figure 5).LL- 37 and citLL-37 significantly enhanced the abundance of Regnase-1 by ≥70% after 30 min, compared to unstimulated PBEC (Figure 5A). Furthermore, the combination of LL-37 or citLL- 37 with IL-17A/F significantly enhanced the abundance of Regnase-1 (>300%), compared to IL- 17A/F alone, after 24 h (Figure 5B). These results demonstrated that both LL-37 and citLL-37 enhances Regnase-1, a RBP known to negatively regulate IL-17-mediated signaling including the production of LCN-2 [23, 44]. As IL-17 signaling and Regnase-1 is known to engage the NF-κB pathway [23, 44, 46], we also examined the abundance of NF-κB p65, and the phosphorylation of p-IKKα/β (S176/180) which is required for the activation of IKK kinases and subsequent NF-κB activation, by Western blots in human PBEC. LL-37, but not citLL-37, significantly enhanced phosphorylation of p-IKKα/β (S176/180) and the abundance of the NF-κB subunit p65 by ∼60%, compared to unstimulated PBEC after 30 mins (Figure 5C).

**Figure 5:**
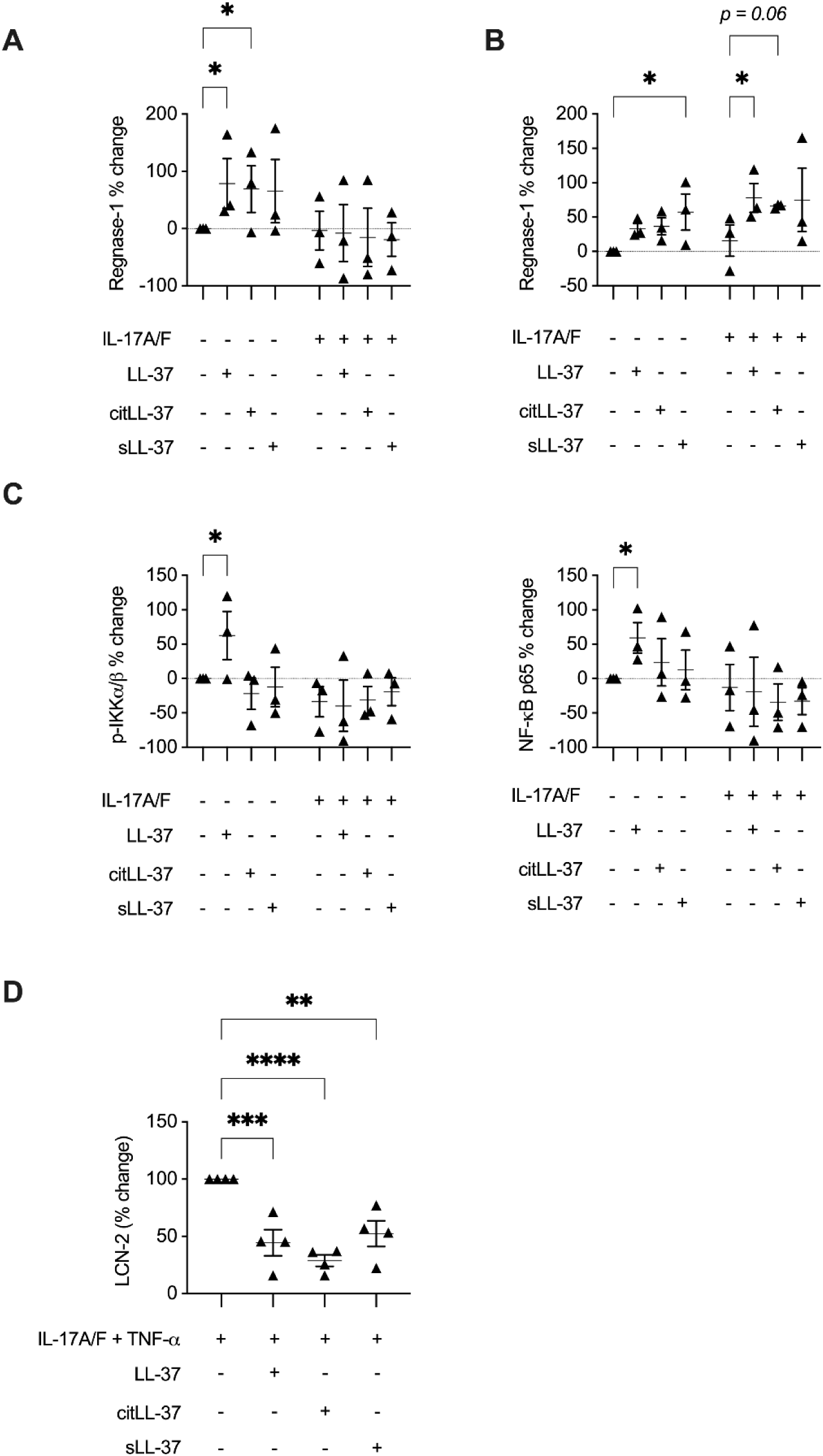
LL-37 and citLL-37 on Regnase-1, p-IKKα/β (S176/180), NF-κB p65 and LCN-2 in human primary bronchial epithelial cells. Human PBEC obtained from three independent donors (N=3) were stimulated with LL-37, citLL-37, or sLL-37 (0.25 μM), in the presence and absence of IL-17A/F (50 ng/mL). Cell lysates (25 μg total protein per sample) was used to examine the abundance of Regnase-1 **(A)** 30 minutes and **(B)** 24 h post-stimulation, by Western blots. **(C)** The abundance of p-IKKα/β (S176/180) and NF-κB p65, 30 minutes post-stimulation, by Western blots. **(D)** Human PBEC obtained from three independent donors (N=3) were stimulated with LL- 37, citLL-37, or sLL-37 (0.25 μM), in the presence and absence of IL-17A/F (50 ng/mL) and TNF-α (20 ng/mL). The abundance of LCN-2, 24 h post-stimulation, by ELISA. Y-axis represents % change compared to paired unstimulated cells from each donor. Each dot represents an independent experiment, and bars show the mean and SEM. Each dot is reported as percent (%) change compared to unstimulated PBEC where % change = ((treatment – control) / control) x 100%. Repeated measures one-way ANOVA with Fisher’s least significant difference test was used for statistical analysis (*\*p≤0.05*).

Finally, we examined the abundance of LCN-2 production by ELISA in PBECs. In these experiments we used the cytomix (IL-17A/F+TNF) as we have demonstrated that the combinatorial effect of these cytokines robustly mediate neutrophil migration, which is mitigated by the immunodepletion of LCN-2 (Figure 3). Cytomix IL-17A/F+TNFα-mediated LCN-2 production was significantly suppressed in the presence of LL-37 and citLL-37 (between 55-70%) in human PBECs after 24 h (Figure 5D). However, in primary cells, sLL-37 also suppressed LCN- 2 production which was not seen in HBEC-3KT cell line. Nevertheless, taken together these results suggest that both LL-37 and citLL-37 enhanced Regnase-1 and suppressed LCN-2 in PBECs. In addition, these results suggest that citrullination of LL-37 may be a post-translational mechanism to limit the pro-inflammatory arm of LL-37 mediated by the selective loss of NF-κB activation, while maintaining the anti-inflammatory functions of the peptide mediated in part post-transcriptionally by engaging the RBP Regnase-1.

### LL-37, but not citLL-37, enhances the production of CCL20 in the presence and absence of IL- 17A/F

Previous studies have demonstrated that IL-17A enhances the production of Th17- recruiting chemokine CCL20 in neutrophilic airway inflammation (48) and in epithelial cells [23, 44]. IL-17A-mediated transcription of CCL20 is solely dependent on IκB-ζ (similar to GROα), and directly targeted by the ribonuclease Regnase-1 for mRNA degradation [44, 45]. Therefore, as an additional line of investigation, we also examined the effect of the peptides on CCL20 production, in the presence/absence of IL-17A/F. HBEC-3KT cells and human PBEC were stimulated with LL-37, citLL-37 or sLL-37 (0.25 μM), in the presence/absence of IL-17A/F (50 ng/mL), and the abundance of CCL20 was measured in TC supernatant by ELISA, 24 h post-stimulation. Only LL-37 significantly enhanced the production of CCL20 in the presence and absence of IL-17A/F-mediated inflammation by ∼121% and ∼190% respectively, compared to unstimulated HBEC-3KT (Figure 6A). In contrast, citLL-37 did not induce CCL20 production, and also abrogated IL-17A/F-mediated CCL20 to baseline levels, in HBEC-3KT cells (Figure 6A). Similar results were also observed in human PBEC. LL-37 significantly enhanced the production of CCL20 in the presence and absence of IL-17A/F by ∼208% and ∼532% respectively, compared to unstimulated PBEC (Figure 6B). In PBECs, although CCL20 was enhanced in response to citLL-37, this was ∼50% less compared to LL-37 alone (Figure 6B). Similarly, although citLL-37 also enhanced IL-17A/F-mediated CCL20 production, this was ∼35% less compared to LL-37, in human PBEC (Figure 6B). Taken together our results indicate that citrullination of LL-37 may be a post-translational homeostatic mechanism to limit inflammation by selectively decreasing levels of chemokines such as CCL20 compared to the unmodified peptide.

**Figure 6:**
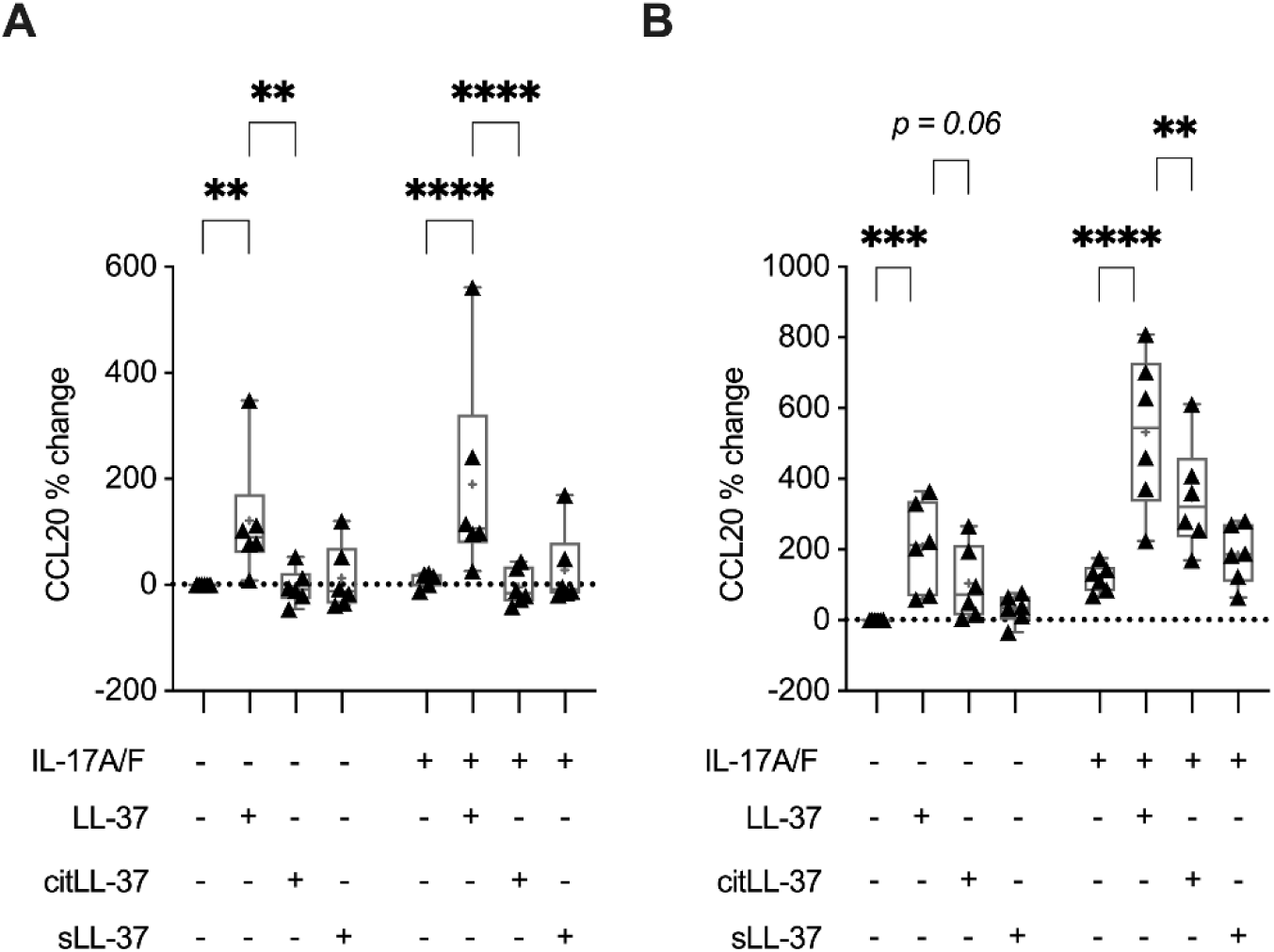
LL-37, but not citLL-37, enhances IL-17A/F-mediated CCL20 production in bronchial epithelial cells. **(A)** HBEC-3KT cells (n=6) and **(B)** human PBEC (N=3 independent donors, n=2 technical replicate each) were stimulated with LL-37, citLL-37 or sLL-37 (0.25 μM), in the presence/absence of IL-17A/F (50 ng/mL), for 24 h. TC supernatants were examined for the abundance of CCL20 by ELISA. Y-axis represents % change compared to paired unstimulated cells for each replicate. Each dot represents an independent experiment, and bars show the median and min-max range. Each dot is reported as percent (%) change compared to unstimulated control bronchial epithelial cells where % change = ((treatment – control) / control) x 100%. Repeated measures one-way analysis of variance with Fisher’s least significant difference test was used for statistical analysis (*\*\*p≤0.001, ***p≤0.005, ****p≤0.0001*).

### Cathelicidin CRAMP, IL-17A/F, LCN-2 and neutrophil elastase (NE) are concurrently increased in a mouse model of neutrophilic airway inflammation

As our *in vitro* results (in bronchial epithelial cells) demonstrated that LL-37 can modulate IL-17-mediated airway inflammation by influencing processes related to neutrophil recruitment, we next examined the abundance of protein targets selected from our *in vitro* studies in a mouse model of neutrophilic airway inflammation [6] (Figure 7A). A previous study has demonstrated in this model (Figure 7A) that a concomitant intranasal (i.n.) administration of HDM and low concentration of LPS during the allergen sensitization phase results in enhanced IL-17-mediated neutrophil accumulation, NET formation, and PADI4-dependent citrullination in the lungs of male mice [6]. In this study, we expanded on this model and report data using both male and female mice. There were no changes to the survival rate (data not shown) or weight of male or female mice after 15 days (Supplemental Figure 6). Leukocyte accumulation was determined by cell differential analysis in BAL, 24 h after the last HDM challenge (Supplemental Figure 6). Male and female mice sensitized with a co-challenge of HDM and LPS showed a clear neutrophil-skewed airway inflammation, compared to mice sensitized with either allergen or endotoxin alone (Figure 7B). Moreover, NE abundance was significantly enhanced in the BALF and lung tissue lysates obtained from mice co-sensitized with HDM and LPS, compared to mice sensitized with either HDM alone or saline controls (Figure 7C). In female mice co-sensitized with HDM+LPS, NE abundance in the BAL and tissue was significantly increased ∼204% and ∼131% respectively, compared to mice sensitized with HDM alone (Figure 7C). In male mice co-sensitized with HDM and LPS, NE abundance in the BALF and tissue was significantly increased ∼77% and ∼82% respectively, compared to those sensitized with HDM alone (Figure 7C). Therefore, mice sensitized with a co-challenge of HDM and LPS resulted in increased accumulation of neutrophils and significant increase in the abundance of NE, a marker of neutrophil activation, in both male and female mice. We further examined the abundance of IL-A/F, cathelicidin CRAMP (mouse analog of LL- 37), and LCN-2 in this mouse model of neutrophilic-skewed airway inflammation. Although patients with chronic lung disease characterized by neutrophil accumulation and elevated NET formation show increased abundance of LL-37 in the lungs, compared to individuals with non-neutrophilic airway inflammation and healthy controls [5], it is unknown whether CRAMP levels are enhanced during IL-17-driven neutrophil accumulation in the lungs in murine models. CRAMP abundance in the BALF of female and male mice co-sensitized with HDM + LPS was significantly increased by ∼146% and ∼197% respectively, compared to HDM alone sensitized mice (Figure 8A). Similarly, IL-17A/F abundance in the BAL of female and male mice co-sensitized with HDM + LPS was significantly increased by ∼71% and ∼61% respectively, compared to HDM alone sensitized mice (Figure 8B). LCN-2 abundance was increased in the BAL and lung tissue lysates of female and male mice co-sensitized with HDM + LPS, compared to either HDM sensitized or saline controls (Figure 8C). However, the increase in LCN-2 levels was modest in BAL compared to that in the lung tissue lysates. In female mice co-sensitized with HDM + LPS, LCN-2 abundance in the BAL was increased ∼8% (*p = 0.06*) but significantly enhanced by ∼61%, compared to HDM alone sensitized mice (Figure 8C). Similarly, in male mice LCN-2 abundance in the BAL and lung tissue lysates was significantly increased ∼12% and ∼91% respectively, compared to HDM alone sensitized mice (Figure 8C). These results showed that there was a concurrent increase CRAMP, IL-17A/F and LCN-2 in mice groups with neutrophilic airway inflammation.

**Figure 7:**
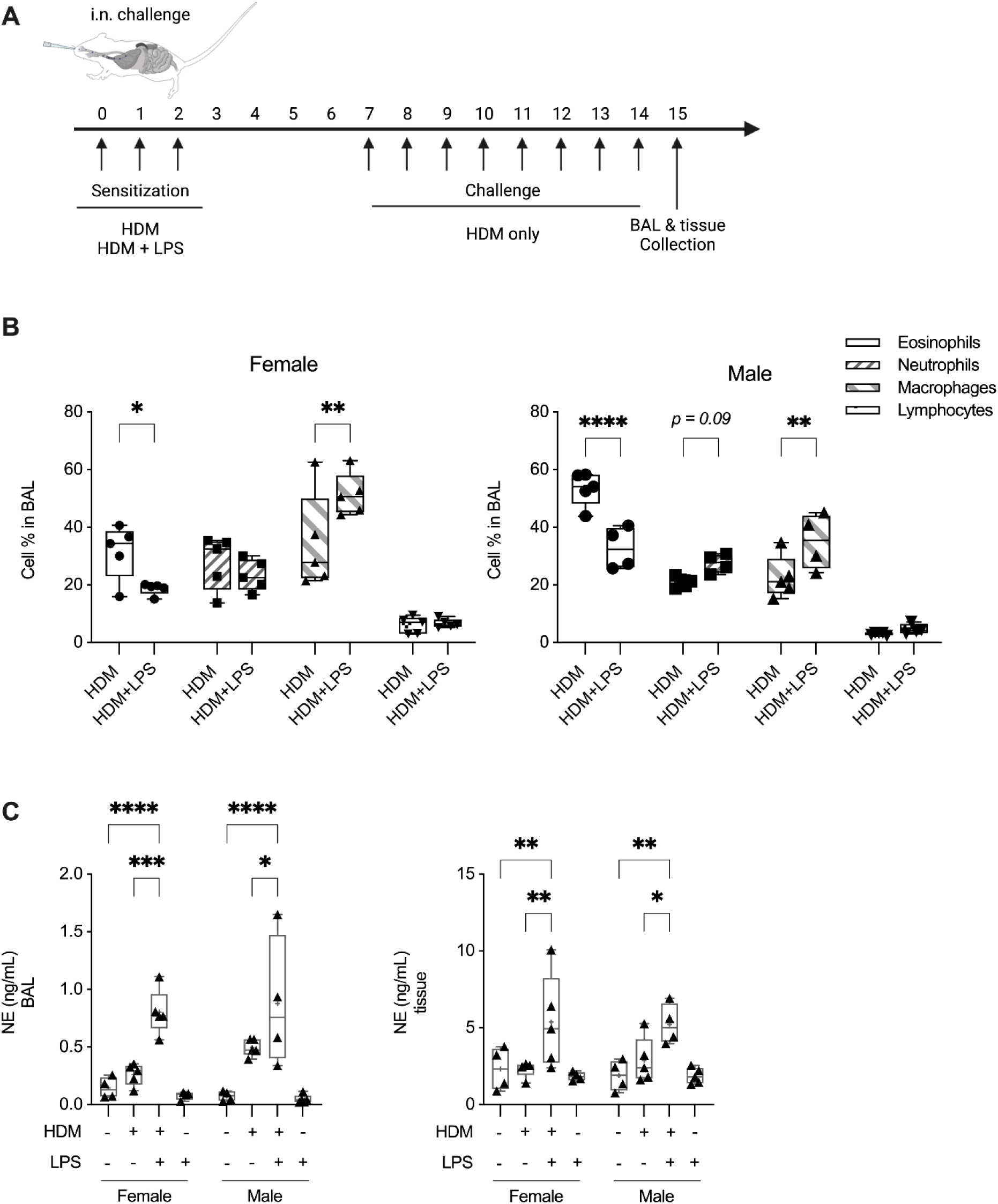
Co-sensitization of mice with allergen house dust mite (HDM) and LPS leads to a neutrophilic skewed airway inflammation. **(A)** Female and male BALB/c mice (8-10 weeks; N≥4 per group) were challenged (i.n.) with either saline, 25 μg of HDM protein extract (35 μL of 7 µg/mL saline per mouse) with or without 1 μg LPS (35 μL of 0.03 µg/mL saline per mouse), or LPS alone, once daily for 3 consecutive days (days 0 to 2), and subsequently rested for 4 days. Beginning on day 7, allergen HDM was administered (allergen recall) in mice (i.n.) sensitized with either HDM alone or with a combination of HDM and LPS, once daily for 8 consecutive days. **(B)** BALF collected from female and male mice 24 h after the last HDM challenge was used for cell differentials, to assess cell percentages ((count of individual cell populations / count total cell accumulation) x 100). **(C)** Abundance of NE in the BALF and lung tissue lysates were assessed by ELISA. Bars show median and IQR, whiskers show minimum and maximum points, + denotes average. Statistical analysis was determined by one-way ANOVA with Fisher’s LSD test (*\*p≤0.05, **p≤0.001, ***p≤0.005, ****p≤0.0001*).

**Figure 8:**
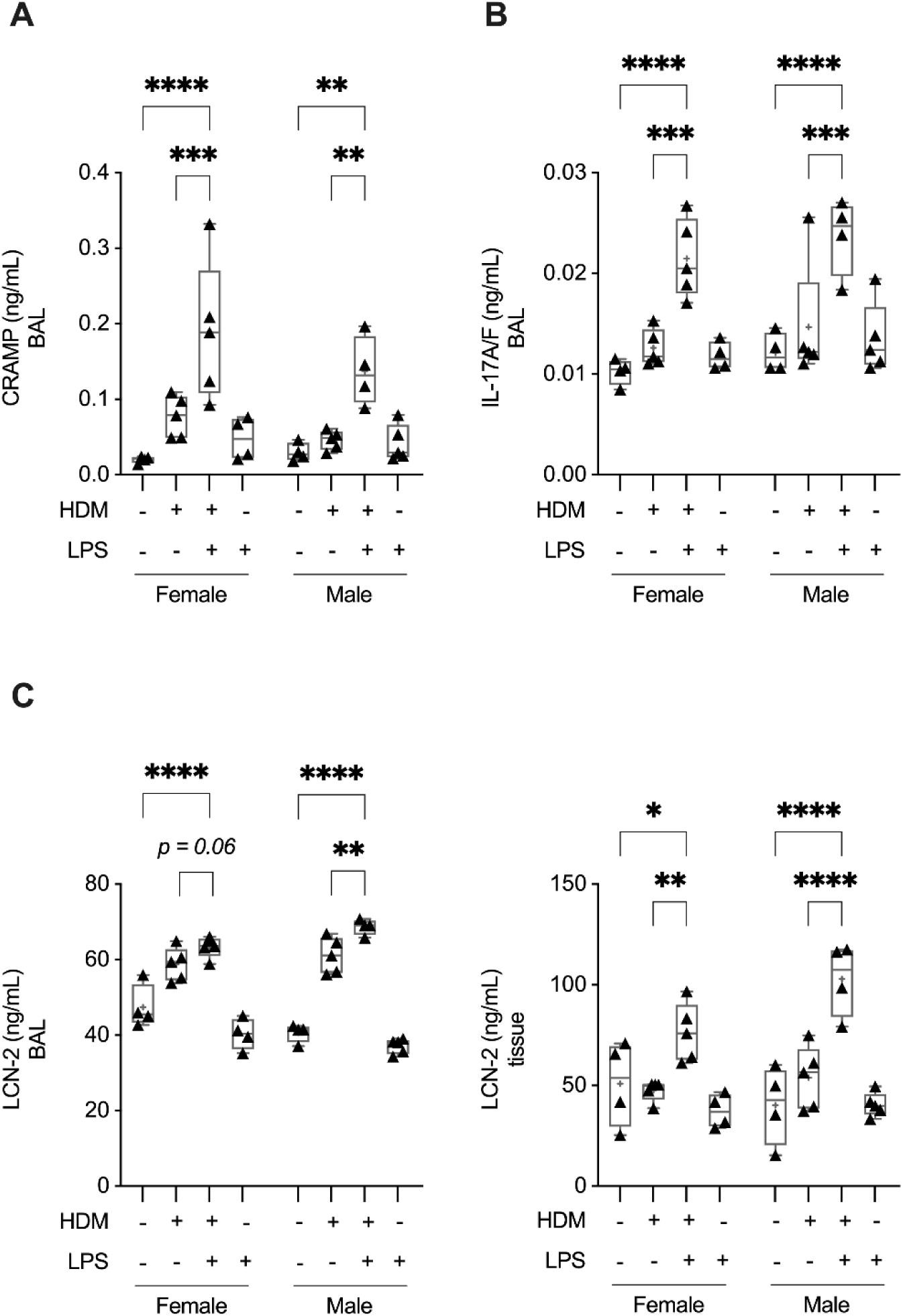
CRAMP, IL-17A/F and LCN-2 are concurrently increased in the lung in the mouse model of neutrophilic airway inflammation. Female and male BALB/c mice (8-10 weeks; N≥4 per group) were challenged (i.n.) with either saline, 25 μg of HDM protein extract (35 μL of 7 µg/mL saline per mouse) with or without 1 μg LPS (35 μL of 0.03 µg/mL saline per mouse), or LPS alone, once daily for 3 days (days 0 to 2), rested for 4 days followed by HDM allergen recall challenge once daily for 8 consecutive days (as detailed in Fig. 8A). Abundance of **(A)** CRAMP and **(B)** IL-17A/F in bronchoalveolar lavage (BAL) was assessed by ELISA, 24 h after the last HDM challenge. **(C)** Abundance of LCN-2 was assessed in the BALF and lung tissue lysates by ELISA. Bars show median and IQR, whiskers show minimum and maximum points, + denotes average. Statistical analysis was determined by one-way ANOVA with Fisher’s LSD test (*\*p≤0.05, **p≤0.001, ***p≤0.005, ****p≤0.0001*). HDM, house dust mite; LPS, lipopolysaccharide.

### CRAMP abundance negatively correlates with LCN-2 and positively correlates with NE in the lungs

To determine the relationship between CRAMP, LCN-2, and neutrophil activity (using NE as a marker) *in vivo*, we performed correlative assessments of the abundance of these targets in the lungs of mice with neutrophilic-skewed airway inflammation (HDM + LPS group), in the mouse model detailed above (Figure 9A). LCN-2 showed a significant positive correlation with the abundance of NE in the lung tissue of female mice (Figure 9A). In contrast, the abundance of LCN-2 in the lung tissue lysates negatively correlated with CRAMP in the BAL of female mice (Figure 9B). In male and female mice, there was a significant negative correlation between CRAMP and neutrophil accumulation in the lungs, however that differed by tissue compartment. In female mice, CRAMP abundance in the lung tissue lysates showed a significant negative correlation with neutrophil accumulation in the lungs, whereas in male mice CRAMP abundance in the BAL showed significant negative correlation with neutrophil accumulation (Supplemental Figure 7).

**Figure 9:**
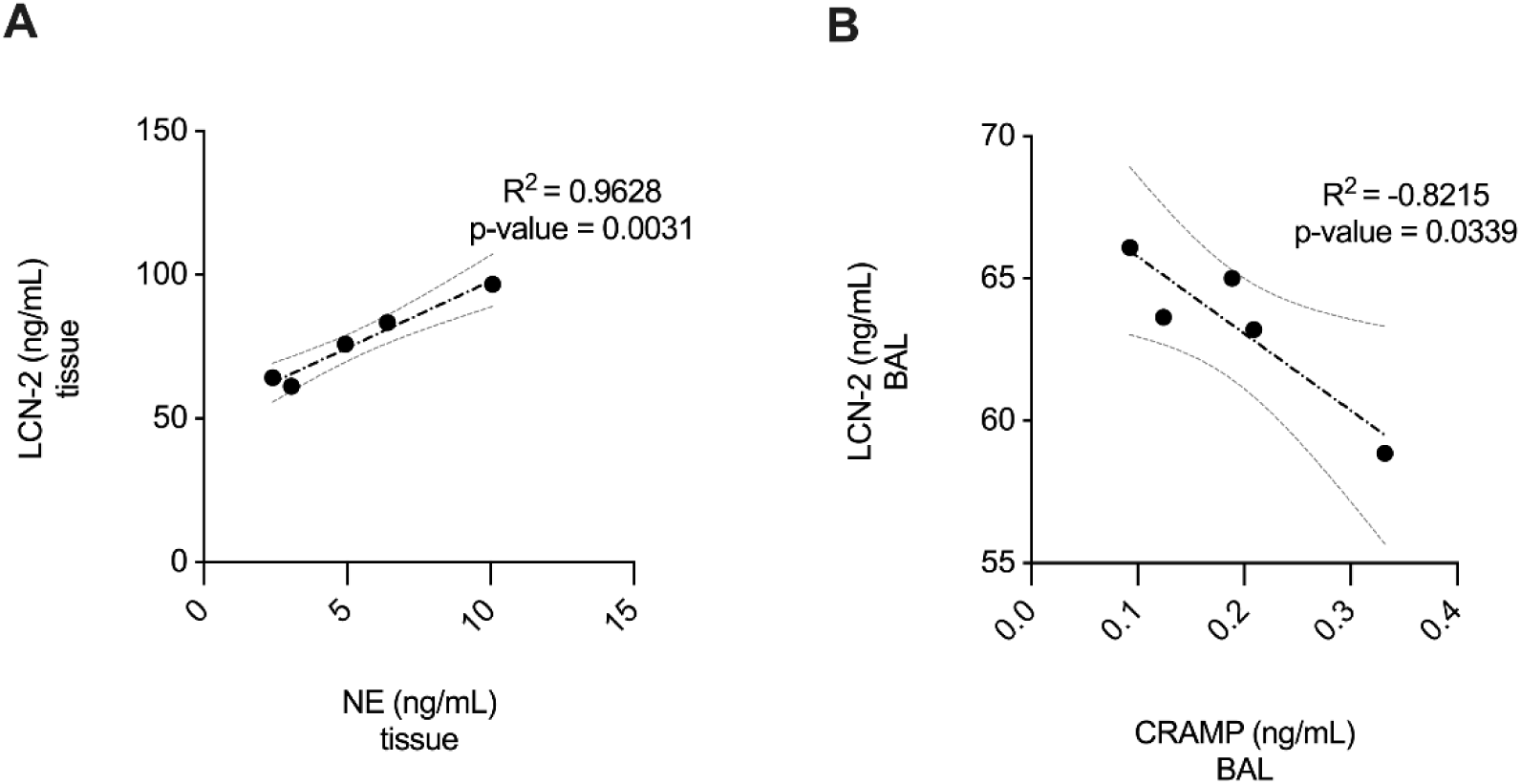
CRAMP abundance shows a negative association with LCN-2, and positive association with NE, in the lungs. Brochoalveolar lavage (BAL) and lung tissue lysates obtained 24 h after the last HDM challenge from mouse model detailed in Figure 7A, were assessed for the abundance of CRAMP and LCN-2 and NE by ELISA. Pearson’s correlation analysis was performed to determine the correlations between **(A)** LCN-2 and NE, and **(B)** LCN-2 and CRAMP, as indicated. *p ≤ 0.05* was considered statistically significant.

## Discussion

In this study, we provide an insight into the interplay of LL-37 and IL-17A/F in the lungs. We identified proteins that are most enhanced in response to IL-17A/F in HBEC, which primarily included neutrophil chemoattractants such as GROα, LCN-2 and Elafin. We provide evidence to support the functional role of LCN-2 in facilitating neutrophil migration, by demonstrating that depletion of LCN-2 within the IL-17A/F-induced inflammatory milieu results in significant decrease in neutrophil migration. We further demonstrated that LL-37 and its citrullinated form, citLL-37, selectively suppressed IL-17A/F-induced LCN-2 abundance in bronchial epithelial cells. This was corroborated by our findings in a mouse model of IL-17-driven neutrophil-skewed airway inflammation [6], wherein we showed a negative correlation between LCN-2 and the mouse cathelicidin peptide CRAMP. Our mechanistic studies suggested that LL-37 and citLL-37 can suppress LCN-2 by (i) suppression of IL-17A/F-mediated C/EBPβ, a transcription factor required for LCN-2 production, and/or (ii) enhancing the ribonuclease regnase-1, which is a negative regulator of IL-17 and LCN-2 [23, 44]. Overall, our findings establish the potential of LL-37 to control neutrophilic airway inflammation by intervening in the production of IL-17-mediated LCN-2. To our knowledge, this is the first study to indicate that the immunomodulatory functions of LL-37 may be mediated by post-transcriptional regulation involving the RBP Regnase-1.

Previous studies have primarily demonstrated negative regulation of cytokine-mediated inflammatory responses by LL-37 primarily in leukocytes. For example, LL-37 suppresses IL-32- induced pro-inflammatory cytokines and enhances the anti-inflammatory cytokine IL-1RA, engaging the SRC Kinase pathway in human macrophages and PBMC [47]. However, LL-37 has been shown to promote cytokine-mediated inflammation in structural cells; LL-37 and IL-17A was shown to induce the transcription of TNFα in human synovial sarcoma cells [19], and LL-37 with IL-1β was demonstrated to synergistically increase IL-8 production in airway epithelial cells [48]. In contrast, the results of this study suggest that LL-37 can enhance certain IL-17A/F- mediated downstream responses such as GROα, while selectively suppressing other responses such as LCN-2, in bronchial epithelial cells. Although both GROα and LCN-2 can facilitate neutrophil recruitment, we demonstrate that depletion of LCN-2 from the secreted inflammatory milieu mitigates neutrophil migration. As LL-37 and citLL-37 can suppress LCN-2 production, our findings suggest that these peptides have the potential to limit neutrophilic airway inflammation.

Interestingly, LCN-2 also functions as an antimicrobial protein in the lungs [49]. Here, we show that an antimicrobial host defence peptide LL-37 can suppress the abundance of another antimicrobial protein, LCN-2, to modulate a cytokine-mediated inflammatory response. This suggests that cathelicidin peptides such as LL-37 can influence the expression and/or activity of other antimicrobial peptides / proteins to regulate the inflammatory cascade. This opens a new avenue of research to examine how the network of various antimicrobial host defence peptides are altered by cathelicidins during inflammation.

Change in the abundance of cathelicidin peptides such as LL-37 seems to be dependent on the kinetic and type of airway inflammation including that with neutrophilia [4, 5, 42, 50, 51]. Moreover, the effect of LL-37 on neutrophils remains confounding; LL-37 can suppress neutrophil chemotaxis by promoting internalization of the chemokine receptor CXCR2 on neutrophils [52], and in contrast directly enhance neutrophil recruitment by activating the formyl peptide receptor 1 [53]. It was previously suggested that LL-37 can enhance GROα in human PBMC [54] and this results in the enhanced recruitment of neutrophils [32]. Interestingly, here we show that despite the presence of GROα in TC supernatants obtained from HBEC, selectively depleting LCN-2 significantly suppresses neutrophil migration. Thus, our results functionally demonstrate the role of LCN-2 in directly facilitating neutrophil migration. This is corroborated by previous studies demonstrating that LCN-2 can enhance neutrophil migration in chronic inflammatory disease [36, 37, 55]. Aligned with this, a previous study showed that recombinant human and mouse LCN-2 enhances neutrophil migration [36]. As we show that LL-37 and citLL-37 can suppress IL-17A/F- mediated LCN-2 production in HBEC, it is likely that that these peptides can limit infiltration of neutrophils in the lungs during airway inflammation.

Neutrophils are a dominant source of cathelicidins in chronic lung disease characterized by airway inflammation [56]. There is a concomitant increase in LL-37 with markers of NET formation such as NE and extracellular DNA [5]. This is aligned with our results demonstrating an increase in the mouse cathelicidin CRAMP along with NE in the lungs, in a mouse model of allergen-challenged airway inflammation [6]. Our *in vivo* results demonstrate that CRAMP, IL- 17A/F, LCN-2 and NE are all concurrently elevated with increased neutrophilic accumulation in the lungs. We show that while LCN-2 positively correlates with NE, its abundance in the lungs negatively correlates with CRAMP (mouse ortholog of LL-37). Thus, our results suggest that cathelicidin peptide secretion by neutrophils in the lungs may be a negative-feedback loop to limit IL-17A/F-induced airway inflammation via peptide-mediated suppression of LCN-2. It is likely that the regulation of IL-17A/F-mediated downstream processes by cathelicidins is dynamic and dependent on the kinetics of response. Cathelicidins can potentiate IL-17A/F-producing Th17 cells in the lung [7], suggesting that LL-37 may drive neutrophil accumulation in the lung during the initiation phase of inflammation. However, it is possible that LL-37 can limit neutrophil infiltration and airway inflammation in later stages of inflammation by intervening in IL-17-mediated LCN- 2 production, such as in chronic airway inflammation.

Our findings indicate that the mechanistic underpinnings of regulation of IL-17A/F-induced downstream responses by LL-37 includes the RBP Regnase-1, which is a inhibitor of IL-17- mediated signaling and LCN-2 production [23, 44]. The immunomodulatory functions of LL-37 involves the peptide’s direct interaction with GAPDH in monocytes [54]. A recent study has shown that GAPDH moonlights as an RBP facilitating degradation of pro-inflammatory cytokine TNFα mRNA and downstream inflammatory response [57]. Similarly, here we demonstrate that LL-37 enhances the RBP Regnase-1, indicating a post-transcriptional mechanism related to LL-37’s ability to selectively alters IL-17A/F-mediated downstream responses. The role of LL-37 in regulating inflammation engaging post-transcriptional machinery is a new area of investigation.

It is known that enzymes PADI2- and PADI4-dependent citrullination of LL-37 occurs in the human lung [20, 42], and that PADI4 activation increases in IL-17-driven neutrophilic airway inflammation [6]. Citrullination of LL-37 impairs the ability of the peptide to limit pathogen or endotoxin-mediated inflammation [20–22, 42, 43]. Citrullination of LL-37 limits endotoxin neutralizing activity of the peptide [21, 42, 43] and increases IL-6 abundance in a model of sepsis [21]. However, here we demonstrate that citrullination does not mitigate all immunomodulatory functions of LL-37. We show that both LL-37 and citLL-37 significantly suppress IL-17A/F- induced LCN-2 abundance and enhances the RBP Regnase-1. In contrast, we show that LL-37, but not citLL37, enhances IL-17A/F-mediated enhancement of *NFKBIZ* mRNA abundance. IκB-ζ (encoded by *NFKBIZ* mRNA) is known to control IL-17A-mediated induction of GROα and CCL20 [23]. Thus, our findings suggest that the differential activity of LL-37 and citLL-37 on IL- 17A/F-mediated GROα and CCL20 production may be due to the selective loss of IκBζ mRNA abundance by citLL-37. Overall, our results indicate that citrullination of LL-37 results in the selective loss of the classical pro-inflammatory NF-κB signal transduction, without altering the increase in ribonuclease Regnase-1 which is an anti-inflammatory mediator. It is thus likely that citrullination of LL-37 may be a mechanism to facilitate immune homeostasis by selectively limiting the peptide’s ‘pro-inflammatory’ functions. To our knowledge, this is the first study to show that citrullination does not impair all immunomodulatory functions of LL-37. Taken together, our results indicate that even though LL-37 may be citrullinated within the IL-17-driven neutrophilic inflammatory milieu, the peptide’s ability to enhance regnase-1 and suppress LCN-2, and consequently dampen neutrophil infiltration in the lungs, is maintained.

A limitation of this study is a lack of direct *in vivo* evidence to support our findings that LL- 37 controls neutrophilic airway inflammation by suppressing LCN-2. This is challenging, as neutrophil recruitment and activation is mitigated in cathelicidin knockout mouse [13], thus making it difficult to directly assess the role of cathelicidin in IL-17-driven neutrophilic airway inflammation *in vivo*. Another limitation is a relatively small sample size in some of the *in vitro* experiments, although we provide statistically significant dataset. Moreover, we have used various complementary approaches to confirm and validate the findings presented in this study. Taken together, the results *in vivo* support our *in vitro* findings, establishing the potential role of LL-37 and citLL-37 in limiting IL-17-driven neutrophilic airway inflammation by selectively intervening in LCN-2 production.

## Conclusion

The findings of this study provide a comprehensive insight into the interplay of LL-37 and IL- 17A/F, relevant to the immunobiology of respiratory disease such as severe asthma and COPD, characterized by airway inflammation with neutrophilia. Our findings indicate that IL-17A/F- mediated increase in LCN-2 may be a central driver of neutrophil migration in the lungs. Our results suggest that cathelicidins such as LL-37 can act as a negative regulator of neutrophil influx to the lungs by selectively suppressing LCN-2. Findings reported in this study support the rationale to examine cathelicidin-derived peptides as interventions to target IL-17-driven neutrophilic airway inflammation for chronic respiratory diseases such as steroid-unresponsive asthma.

## DECLARATIONS

### Ethics approval

The animal model protocol used in this study was approved by the University of Manitoba Animal Research Ethics Board (protocol #18-038). Anonymized patients undergoing resection surgery at the Leiden University Medical Center (LUMC), the Netherlands, were enrolled with a no-objection system for coded anonymous further use of such tissue (www.coreon.org). Since 01-09-2022, patients are enrolled with written informed consent in accordance with local regulations from the LUMC biobank with approval by the institutional medical ethical committee (B20.042/Ab/ab and B20.042/Kb/kb).

### Availability of data and materials

All data generated and analyzed during this study are included in the published article and supplementary information.

### Author Contributions

NM and AA conceived and designed the study. AA performed majority of the experiments, analyzed all the data, and wrote the manuscript. DL assisted with Western blot assays. PR and MH performed the experiments with primary human bronchial epithelial cells. MH performed the animal model study and provided significant intellectual input into study design. AMD provided the human primary bronchial cells, intellectual input for the study, and edited the manuscript. NM obtained funding and resources for the study, provided overall supervision, and edited the manuscript. All the authors reviewed the manuscript.

## Acknowledgements

Not applicable.

