## Supplemental Information for "LL-37 modulates IL-17A/F-mediated airway inflammation by selectively suppressing Lipocalin-2"

Dr. Neeloffer Mookherjee

**Short Title:** LL-37 modulates IL-17-mediated airway inflammation

**FIGURES**

**Supplemental Figure 1:** ***Physiological concentration of LL-37 enhances GRO$\alpha$ and IL-8 secretion.*** HBEC-3KT cells were stimulated with either LL-37, or sLL-37 at different concentrations as indicated, for 24 h. Abundance of chemokines GROα and IL-8 was monitored in the TC supernatants by ELISA. Each dot represents an independent experiment compared to paired unstimulated HBEC-3KT, and dashed lines show the average. Each dot represents an independent experiment, and bars show the median and min-max range. Repeated measures one-way ANOVA with Fisher's least significant difference test was used for statistical analysis (**p≤0.05, **p≤0.001, ***p≤0.005, ****p≤0.0001*).

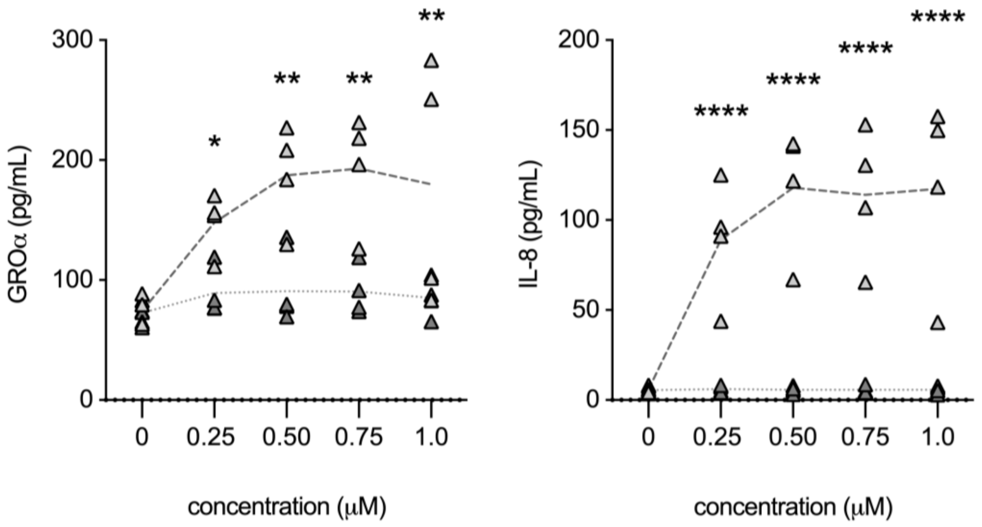

**Supplemental Figure 2:** ***LL-37 and citLL-37 differentially enhance neutrophil chemokines.*** HBEC-3KT cells were stimulated with either LL-37, citLL-37 or sLL-37 (0.25 μM) for 24 h. TC supernatants were examined for the abundance of GROα and IL-8 by ELISA. Each dot represents an independent experiment, and bars show the median and min-max range. Repeated measures one-way ANOVA with Fisher's least significant difference test was used for statistical analysis (**p≤0.05, **p≤0.001, ***p≤0.005*).

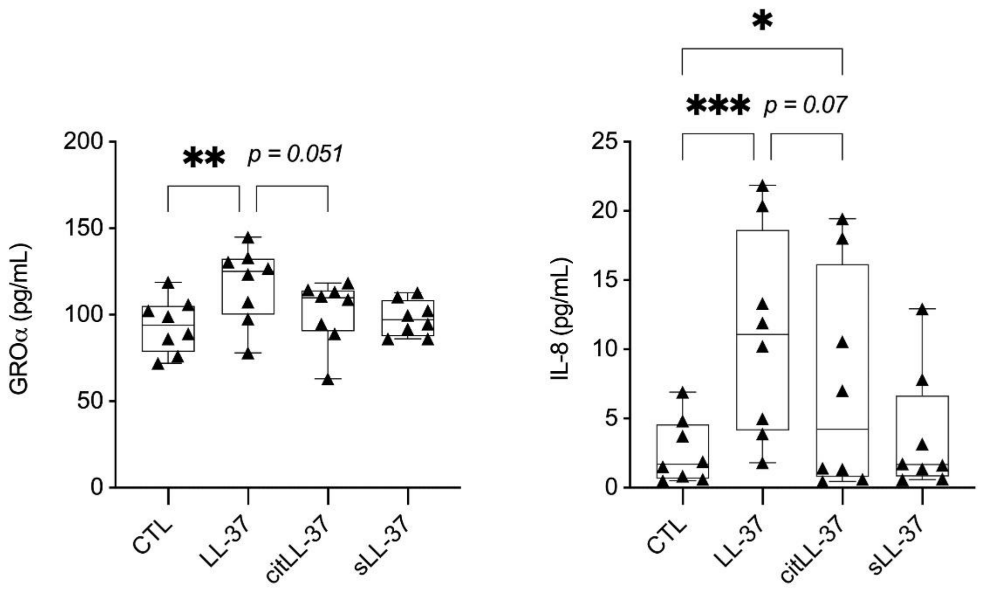

**Supplemental Figure 3:** ***Antibody-mediated neutralization of LCN-2 in TC supernatant.*** HBEC-3KT were stimulated with IL-17A/F (50 ng/mL) and TNFα (20 ng/mL) as indicated. TC supernatants were collected 24 h post-stimulation and incubated with either anti-LCN-2 or anti-IgG antibodies, following which the abundance of LCN-2 was examined by ELISA. Each dot represents an independent experiment, and bars show the median and min-max range. Repeated measures one-way ANOVA with Fisher's least significant difference test was used for statistical analysis (***$p \leq 0.01$*).

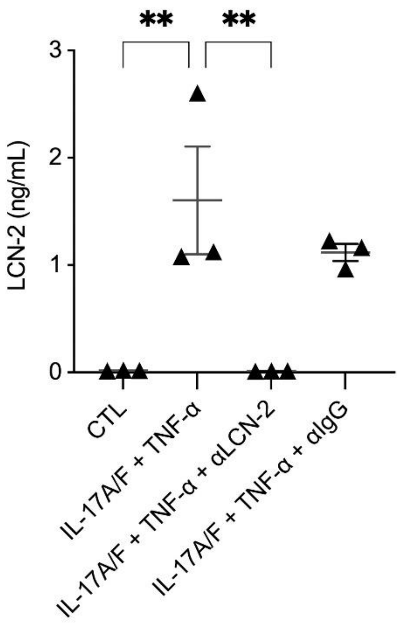

**Supplemental Figure 4:** ***LL-37 and citLL-37 modulates Arid5a and Regnase-1 abundance.*** HBEC-3KT (N=3) were stimulated with IL-17A/F (50 ng/mL) or the peptides LL-37, citLL-37, and sLL-37 (0.25 μM). Total cell lysate (25 μg per sample) was used to determine the abundance of Arid5a and Regnase-1 by Western blot, after **(A)** 30 mins and **(B)** 24 h stimulation. Y-axis represents percent (%) change compared to paired unstimulated cells (controls). Each dot represents an independent experiment, and bars show the mean and SEM. Each dot is reported as % change compared to unstimulated control, where % change = ((treatment – control) / control) x 100%. Repeated measures one-way ANOVA with Fisher's least significant difference test was used for statistical analysis (*$*p \leq 0.05$, $**p \leq 0.001$, $****p \leq 0.0001$*). **(C)** Representative Western blot image, using actin for protein loading control and normalization of densitometry.

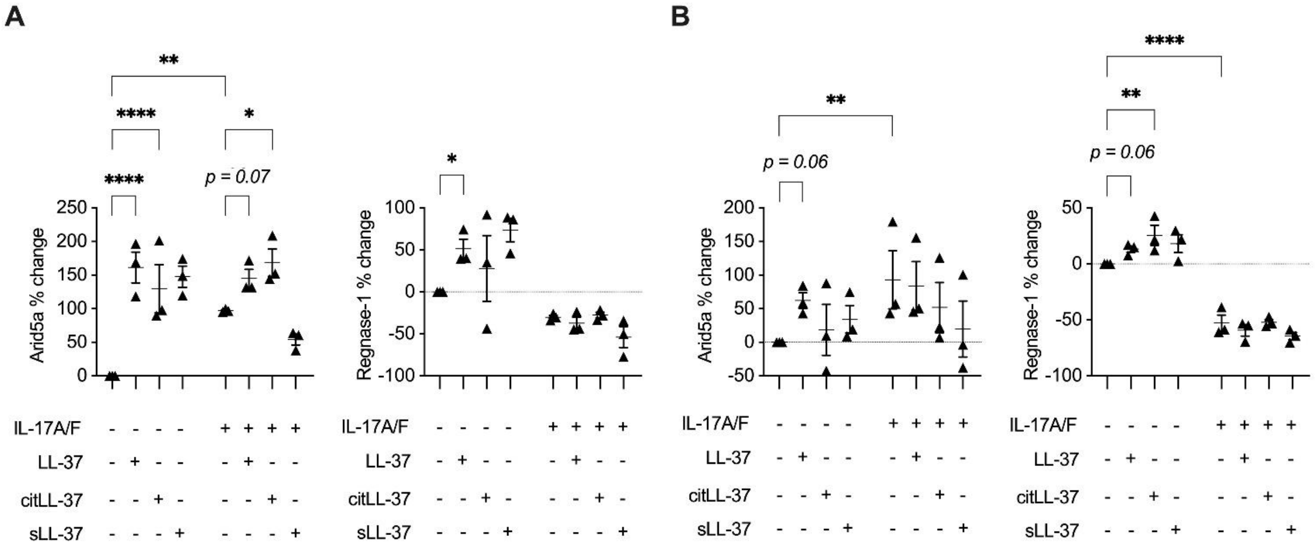

**C**

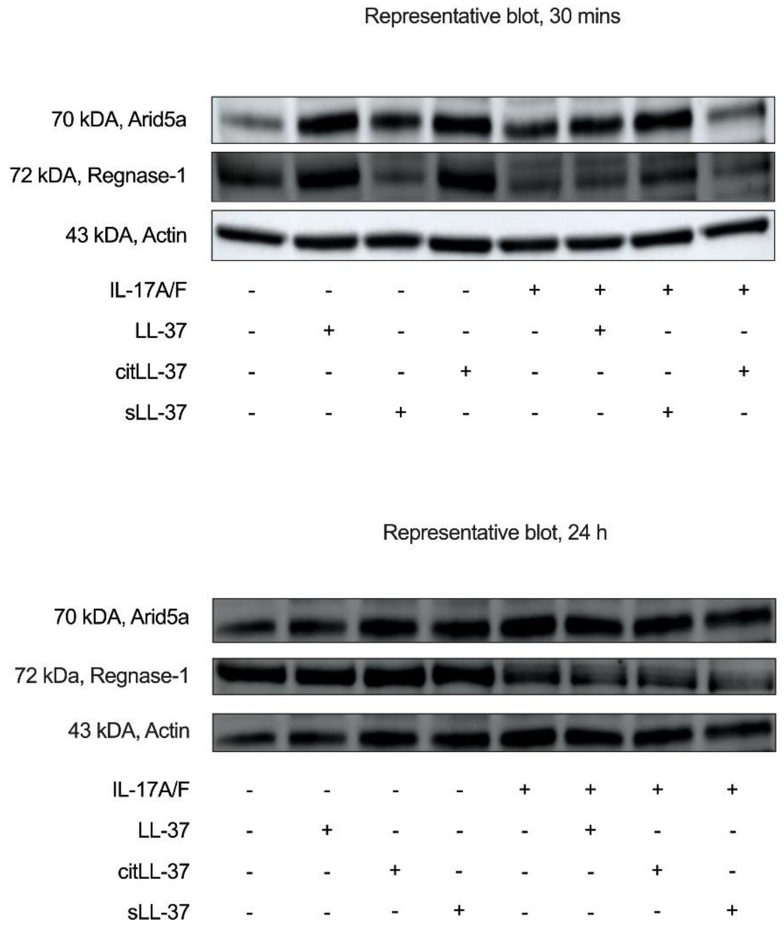
Representative blot, 30 mins
70 kDA, Arid5a
72 kDA, Regnase-1
43 kDA, Actin
IL-17A/F - - - - + + + +
LL-37 - + - - - + - -
citLL-37 - - - + - - - +
sLL-37 - - + - - - + -
Representative blot, 24 h
70 kDA, Arid5a
72 kDa, Regnase-1
43 kDA, Actin
IL-17A/F - - - - + + + +
LL-37 - + - - - + - -
citLL-37 - - + - - - + -
sLL-37 - - - + - - - +

**Supplemental Figure 5:** ***Representative Western blots for human primary bronchial epithelial cells (PBEC).*** Human PBEC from three independent donors (N=3) were stimulated with LL-37, citLL-37, or sLL-37 (0.25 μM), in the presence and absence of IL-17A/F (50 ng/mL). Cell lysates (25 μg total protein per sample) was used to examine the abundance of p-IKKα/β, NF-κB p65 and Regnase-1, after 30 minutes and 24 h, by Western blots. Actin was used to as protein loading control for normalization of densitometry data. Figure shown is a representative of the Western blots using PBEC.

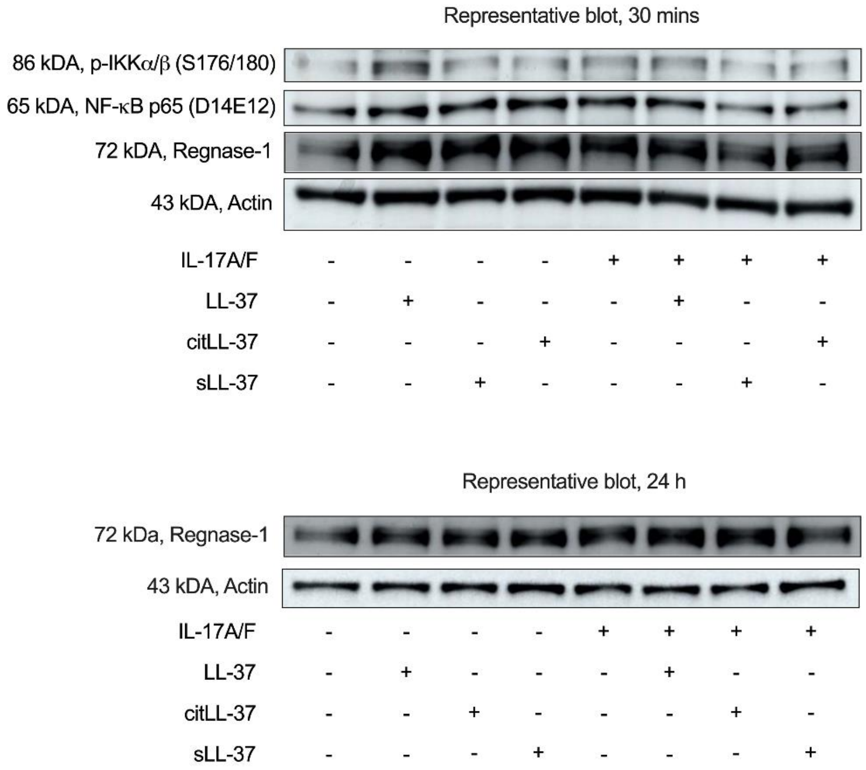

**Supplemental Figure 6:** ***Immune cell accumulation in the lungs of female and male mice, in a mouse model of HDM and LPS challenge.*** Female and male BALB/c mice (8-10 weeks; N≥4 per group) were challenged (i.n.) with either saline, 25 µg of HDM protein extract (35 µL of 7 µg/mL saline per mouse) with or without 1 µg LPS (35 µL of 0.03 µg/mL saline per mouse), or LPS alone, once daily for 3 consecutive days (days 0 to 2), and subsequently rested for 4 days. Beginning on day 7, allergen HDM was administered (allergen recall) in mice (i.n.) sensitized with either HDM alone or with a combination of HDM and LPS, once daily for 8 consecutive days. **(A)** Bodyweight measurement. Bronchoalveolar lavage fluid was collected 24 h after the last HDM challenge and used to assess **(B)** neutrophils, **(C)** eosinophils, **(D)** macrophage, and (**E**) lymphocytes. Bars show median and IQR, whiskers show minimum and maximum points, + denotes average. Statistical analysis was determined by one-way ANOVA with Fisher's LSD test (*$*p \leq 0.05$, $**p \leq 0.001$, $***p \leq 0.005$, $****p \leq 0.0001$*). HDM, house dust mite; LPS, lipopolysaccharide.

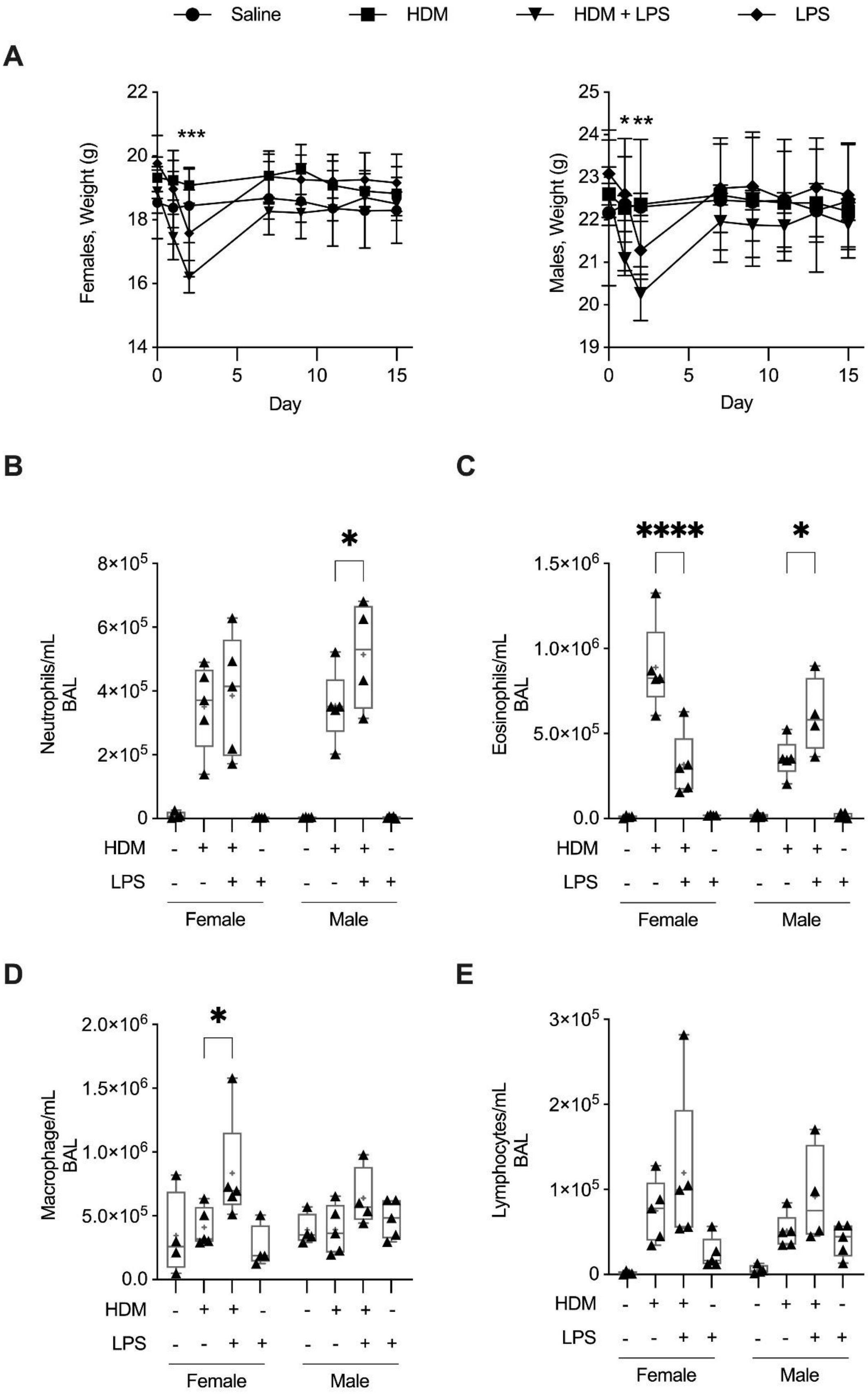
Saline
HDM
HDM + LPS
LPS
A
Females, Weight (g)
Males, Weight (g)
Day
***
B
Neutrophils/mL BAL
C
Eosinophils/mL BAL
D
Macrophage/mL BAL
E
Lymphocytes/mL BAL
HDM
LPS
Female
Male

**Supplemental Figure 7:** ***CRAMP is negatively correlated with neutrophil accumulation in the lungs of mice.*** BALB/c mice (8-10 weeks; N≥4 per group) were challenged (i.n.) with either saline, 25 µg of HDM protein extract (35 µL of 7 µg/mL saline per mouse) with or without 1 µg LPS (35 µL of 0.03 µg/mL saline per mouse), or LPS alone, once daily for 3 consecutive days (days 0 to 2), and subsequently rested for 4 days. Beginning on day 7, allergen HDM was administered (allergen recall) in mice (i.n.) sensitized with either HDM alone or with a combination of HDM and LPS, once daily for 8 consecutive days. Bronchoalveolar lavage (BAL) and lung tissue were collected 24 h after the last HDM challenge. Abundance of CRAMP was assessed by ELISA. Neutrophil numbers in BAL were assessed by cell differential assessment. Pearson's correlation analysis was performed to determine the correlations between CRAMP and neutrophils. Figures shown are **(A)** lung tissue lysates from female mice, and **(B)** BAL from male mice. *$*p < 0.05$* were considered statistically significant.

**(A)** **(B)**

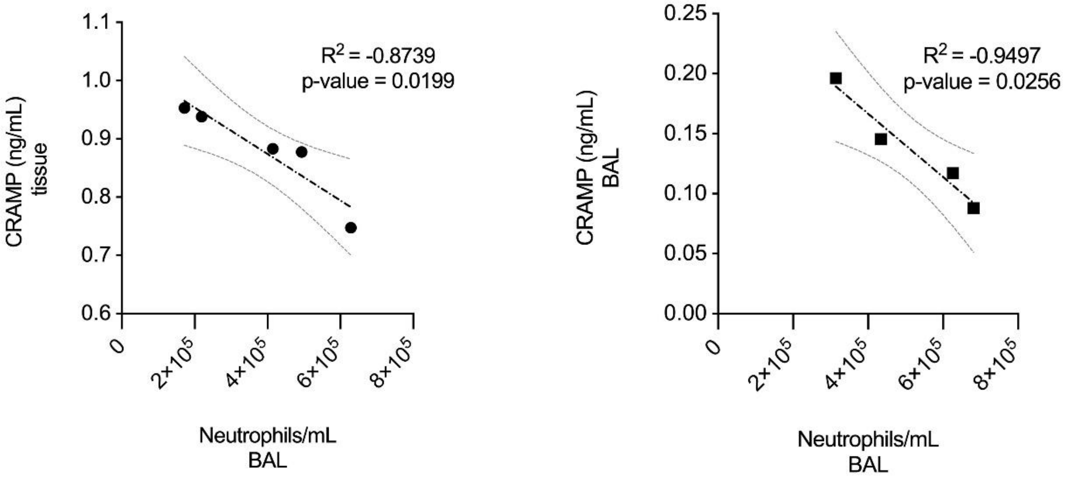

**Supplemental Table I: Proteins significantly altered in response to IL-17A/F in human bronchial epithelial cells.**

| NAME | SWPROT ID | AVG LOG2 FC (normalized to unstimulated cells) | p-value |
|---|---|---|---|
| **LCN-2** | **P80188** | **1.86** | **0.00039384** |
| **ELAFIN** | **P19957** | **1.64** | **0.00735901** |
| STC1 | P52823 | 1.27 | 0.00233145 |
| IGFBP5 | P24593 | 0.74 | 0.00638196 |
| IGFBP3 | P17936 | 0.57 | 0.00283743 |
| IL17A | Q16552 | 0.49 | 0.02162217 |
| SLPI | P03973 | 0.41 | 0.01828189 |
| FLT3 | P36888 | 0.25 | 0.04376196 |
| KIR2DL4 | Q99706 | 0.19 | 0.04528727 |
| SAA1 | P0DJI8 | 0.17 | 0.00725448 |
| **CXCL1 (GROα)** | **P09341** | **0.12** | **0.00062497** |
| EDA2R | Q9HAV5 | 0.08 | 0.04865734 |
| CXCL13 | O43927 | 0.08 | 0.03957933 |
| LEPR | P48357 | 0.08 | 0.00857008 |
| TNFSF13B | Q9Y275 | 0.07 | 0.02355169 |
| PDGFRA | P16234 | 0.06 | 0.04654336 |
| IBSP | P21815 | 0.05 | 0.0190153 |
| BMPR1A | P36894 | 0.04 | 0.04357518 |
| OMD | Q99983 | 0.04 | 0.03799716 |
| UNC5C | O95185 | 0.03 | 0.0221567 |
| VEGFA | P15692 | -0.07 | 0.02045688 |
| LGALS3BP | Q08380 | -0.17 | 0.03782374 |
| CST3 | P01034 | -0.23 | 0.03730624 |
| NRP1 | O14786 | -0.3 | 0.02429623 |
| CTSV | O60911 | -0.57 | 0.04598988 |
